# Intrabody-guided synapse proteomics defines pyramidal neuron input architecture and uncovers early remodeling in a mouse model of Alzheimer’s disease

**DOI:** 10.1101/2025.11.24.689247

**Authors:** Dan Dascenco, Gabriele Marcassa, Anastasia Mallopoulou, Jeroen Vandensteen, Lars Lefever, Elke Leysen, Blanca Lorente-Echeverría, Artemis Koumoundourou, Danie Daaboul, Bart Ghesquière, Pedro Magalhães, Joris de Wit

## Abstract

Across their proximal and distal dendritic domains, pyramidal neurons (PNs) integrate inputs that differ in morphology and function. Hippocampal CA1 PNs are among the earliest affected neurons in Alzheimer’s disease (AD), but the molecular composition of their inputs and selective vulnerability remain poorly defined. We develop an intrabody-guided proximity-labeling strategy that targets the biotin ligase TurboID to endogenous postsynaptic scaffolds for cell-autonomous mapping of postsynaptic proteomes. Targeting PSD95 or Homer1 enables selective labeling of excitatory postsynaptic proteins in mouse CA1 PNs and resolves subsynaptic organization by comparing the two probes. Mapping the proteomes of major CA1 inputs uncovers a proximal-distal molecular logic that underlies their distinct properties. Applying this approach in the *App^NL-G-F^* AD mouse model reveals an early signaling-driven phase of synaptic remodeling followed by a later translation-linked phase, with persistent downregulation of glutamatergic components. These results provide a molecular atlas of CA1 PN inputs and identify stage-specific mechanisms of synaptic vulnerability in early AD.

## Introduction

Pyramidal neurons (PNs), such as those found in the cerebral cortex and hippocampus, are characterized by large, complex dendritic trees organized into proximal and distal domains that receive inputs from distinct presynaptic partners. This dendritic domain organization plays a critical role in the integration of synaptic inputs by PNs (Magee, 1998; Spruston, 2008; Larkum *et al*., 2009). To compensate for their greater electrotonic distance from the soma and reduced influence on action potential generation, distal inputs have structural and functional adaptations compared to proximal inputs. For instance, the distal inputs of hippocampal CA1 neurons exhibit a larger proportion of perforated and shaft synapses and a higher NMDA receptor/AMPA receptor (NMDAR/AMPAR) ratio (Megías *et al*., 2001; Otmakhova, Otmakhov and Lisman, 2002; Nicholson *et al*., 2006). While the structural and physiological organization of proximal and distal inputs has been described, little is known about the molecular architecture of PN synapses that underlies these input-specific features.

The proximal dendritic domain of hippocampal CA1 PNs receives local input from CA3 PNs via the Schaffer collaterals (SCs), whereas the distal domain receives input from the entorhinal cortex via the temporoammonic (TA) pathway. CA1 PNs integrate these convergent hippocampal and cortical information streams to generate output that is essential for episodic memory. Their selective vulnerability in Alzheimer’s disease (AD), in which CA1 neurons are among the earliest affected neuronal populations, correlates with memory impairments in the symptomatic stages of the disease. Characterizing the molecular architecture of CA1 PN inputs, and how this might change in the presymptomatic stages of AD, has remained challenging due to a lack of tools capable of mapping synaptic protein composition in a cell type-specific manner.

Profiling the protein composition of CA1 PN inputs across their dendritic trees requires synapse-targeted proteomic strategies (Schreiner *et al*., 2017; van Oostrum and Schuman, 2025). An ideal approach would achieve both cell type- and input type-specificity while enabling deep proteomic coverage and minimizing perturbation of synaptic protein composition (Marcassa, Dascenco and de Wit, 2023). Classic biochemical approaches such as subcellular fractionation and affinity purification have provided insight into synaptic organization but typically require large amounts of tissue, average across cell types, and lack spatial precision. Fluorescence-assisted synaptosome sorting (FASS) has expanded access to protein composition of specific synapse types across the brain (Biesemann *et al*., 2014; Apóstolo *et al*., 2020; Paget-Blanc *et al*., 2022; van Oostrum *et al*., 2023; Kaulich *et al*., 2025). However, because synaptosomes contain mixtures of pre- and postsynaptic elements, as well as glial components, the cell type-specificity of this approach is limited. Proximity labeling offers a powerful alternative, enabling proteomic interrogation within specific cell types via engineered biotin ligases (Roux *et al*., 2012; Lam *et al*., 2014; Branon *et al*., 2018). Current implementations of proximity labeling generally rely on overexpressed fusion proteins to target biotin ligases to synapses or do not achieve input-specific resolution (Uezu *et al*., 2016; Rayaprolu *et al*., 2022; Marcassa *et al*., 2025).

Here, we develop an intrabody-based proximity labeling strategy to map the molecular architecture of mouse CA1 PN inputs. By fusing the biotin ligase TurboID to intracellular binders (intrabodies) directed against postsynaptic scaffold proteins, we achieve minimally invasive labeling of postsynaptic proteins in CA1 PNs. We demonstrate that this approach yields spatially accurate representations of synaptic protein organization within the postsynaptic density (PSD) of CA1 PN excitatory inputs. Using this method to map the molecular architecture of the major CA1 inputs, we identify input-specific assemblies of glutamate receptors, auxiliary proteins and ion channels, and uncover a trans-synaptic molecular scaffold that underlies this organization along the proximal-distal dendritic axis. Profiling CA1 excitatory inputs in *App^NL-G-F^* mice, a model of amyloid-beta (Aβ) pathology in AD (Saito *et al*., 2014), reveals early remodeling of synaptic composition, including dysregulation of glutamate receptors, molecular scaffolds, signaling components and trafficking machinery.

Together, these findings provide deep, high-resolution mapping of the molecular composition of CA1 PN inputs within intact hippocampal circuits and uncover stage-specific perturbations of synaptic architectures in early AD. Intrabody-based targeting of TurboID is broadly adaptable to other neuron types and brain regions, and the modularity of the approach enables extension to additional subcellular compartments. Beyond characterizing the molecular composition of synapses, these datasets provide a foundation for more realistic modeling of neuronal signal integration in CA1 PNs.

## Results

### Efficient biotinylation of postsynaptic proteins using intrabody-TurboID

TurboID is a promiscuous biotin ligase that covalently labels neighboring proteins for subsequent identification by streptavidin-mediated affinity purification followed by mass spectrometry (MS) analysis (Figure 1A, (Roux *et al*., 2012; Branon *et al*., 2018)). To target TurboID to postsynapses, we coupled it to intrabodies, engineered protein binders that target endogenous proteins (Gross *et al*., 2013; Holzer *et al*., 2022). To match their expression level with that of their target, intrabody-TurboID expression is regulated by a transcriptional feedback loop in which non-bound fusion proteins repress expression levels (Figure 1B, (Gross *et al*., 2013)).

**Figure 1.**
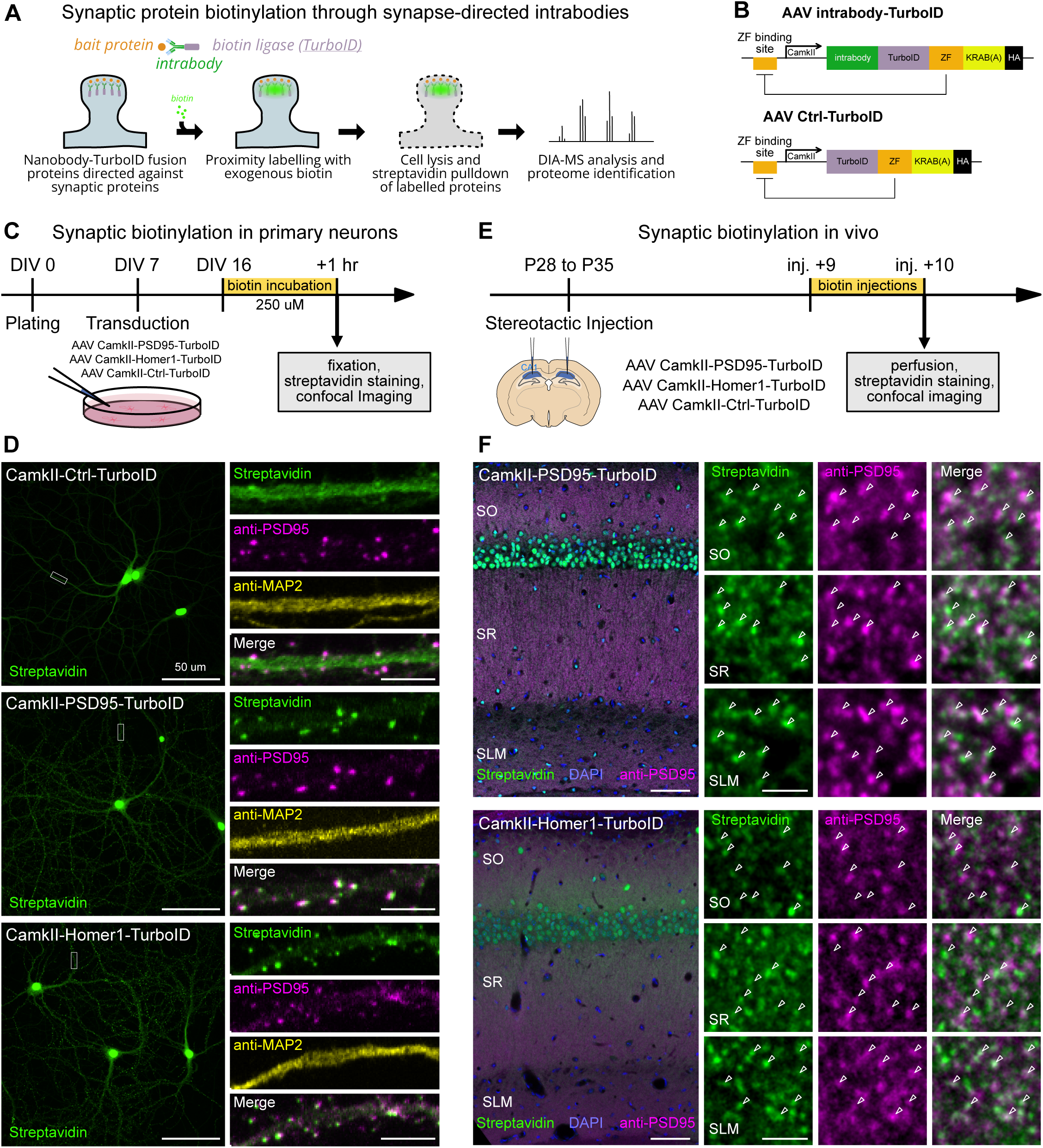
PSDG5-TurboID and Homer1-TurboID efficiently biotinylate excitatory synaptic proteomes in primary cultures and *in vivo.* A. Schematic of intrabody-TurboID-mediated synaptic protein biotinylation. B. Schematic of the AAV constructs used in this study. C. Workflow of intrabody-TurboID validation in primary cultures. D. Validation of the approach in primary cultures shows the localization of biotinylated proteins (probed using Streptavidin-488) for Ctrl-TurboID, PSD95-TurboID and Homer1-TurboID. Left: low magnification z-stacks showing whole neurons, note the nuclear signal. Right: High-magnification z-stack showing cytosolic (Ctrl-TurboID) and synaptic (PSD95-TurboID and Homer1-TurboID) biotinylation, as evidenced by staining against endogenous PSD95 and a dendritic marker MAP2. E. Workflow of intrabody-TurboID validation *in vivo* in hippocampal CA1. F. Validation of the approach in hippocampal CA1. Left: Low-magnification z-stacks of dorsal CA1 in animals injected with PSD95-TurboID and Homer1-TurboID. Right: High-magnification z-stacks showing synaptic biotinylation *in vivo* in the three CA1 synaptic laminae. Scale bars: 50 μm (D left), 5 μm (D right), 100 μm (F left), 2 μm (F right).

To achieve efficient labeling of postsynaptic proteins, we selected two intrabodies directed against excitatory PSD scaffold proteins: a fibronectin type III (FNIII)-based monobody targeting PSD95 (Xph20 (Rimbault *et al*., 2024)), and a camelid nanobody against Homer1 (HC87 (Dong *et al*., 2019)). The regulatory feedback loop drives strong nuclear localization of intrabody fusions, in addition to their desired localization (Gross *et al*., 2013). To exclude off-target nuclear proteins from proteomic identification, we designed a cytosolic control consisting solely of the TurboID-repressor fusion protein (Ctrl-TurboID, Figure 1B).

We generated HA-tagged intrabody-TurboID constructs targeting PSD95 and Homer1 (hereafter termed PSD95-TurboID and Homer1-TurboID), as well as the Ctrl-TurboID construct, downstream of the neuron-specific CamkII promoter (Figures 1B, S1A), and transduced mouse primary neurons using adeno-associated virus (AAV). Following biotin incubation, all constructs exhibited nuclear biotinylation due to the feedback loop, but while the Ctrl AAV displayed diffuse biotinylation in dendrites, both PSD95-TurboID and Homer1-TurboID showed strong punctate labeling colocalizing with the excitatory postsynaptic marker PSD95 (Figure 1D).

We next assessed *in vivo* synaptic labeling by injecting intrabody-TurboID AAVs in the hippocampal CA1 region of wild-type (WT) mice (Figure 1E). Following systemic biotin delivery, both PSD95- and Homer1-TurboID probes displayed a punctate pattern that co-localized with PSD95, in addition to the nuclear localization also seen in cultures (Figure 1F). Biotinylation of postsynaptic proteins extended across the entire length of the CA1 dendritic tree spanning *stratum oriens* (SO), *stratum radiatum* (SR) and *stratum lacunosum-moleculare* (SLM) (Figure 1F). In contrast, the Ctrl-TurboID probe retained the nuclear signal but did not show synaptic biotinylation (Figure S1B).

Together, these results show that the generated intrabody-TurboID probes efficiently biotinylate postsynaptic excitatory proteins in cultured neurons and in CA1 PNs.

### Profiling CA1 PN excitatory postsynaptic proteomes with subsynaptic specificity

To analyze protein composition of CA1 PN inputs, we injected PSD95-, Homer1- and Ctrl-TurboID AAVs in the hippocampal CA1 region and quantified labeling efficiency and proteomic depth of PSD95-TurboID and Homer1-TurboID probes (Figure 2A). Together with the Ctrl-TurboID group, a non-injected group (no_virus) serves to exclude endogenously biotinylated proteins, contaminants and nuclear labelled proteins from the analysis. Western blot (WB) analysis of micro-dissected CA1 tissue extract confirmed probe expression and biotinylation efficiency (Figure 2B). Expression and biotinylation levels were higher for PSD95- and Homer1-TurboID than for Ctrl-TurboID, which predominantly localizes to the nucleus.

**Figure 2.**
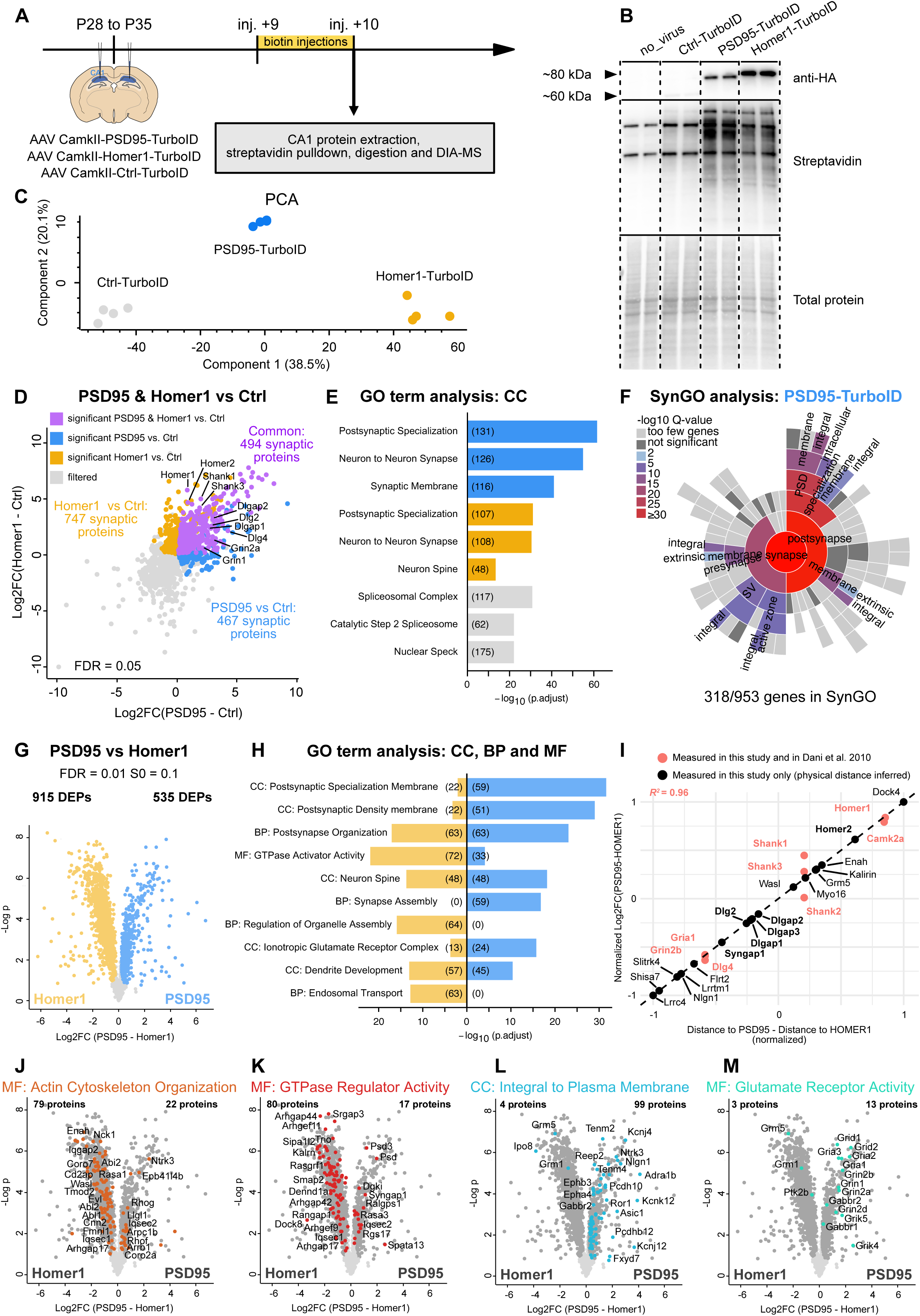
PSDG5-TurboID and Homer1-TurboID target spatially resolved subsynaptic regions. A. Experimental workflow (n=4 replicates per condition, 2/3 animals per replicate, a mix of males and females). B. Western Blot showing expression of intrabody-TurboID and Ctrl-TurboID probes (anti-HA), as well as biotinylation levels (streptavidin-HRP for two out of four replicates in the experiment). C. PCA showing separation of different replicates based on MS intensities. D. Filtering against the Ctrl-TurboID condition shows the number of common and different proteins significantly enriched in the PSD95-TurboID and Homer1-TurboID conditions (ANOVA with Benjamini-Hochberg FDR=0.05 followed by Tukey’s HSD). E. Top 3 GO:CC terms enriched in Homer1-TurboID, PSD95-TurboID and filtered proteomes. Numbers in parenthesis indicate the amounts of annotated proteins in each term. F. SynGO enrichment diagram for the PSD95-TurboID condition. G. Volcano plot showing the enrichment of synaptic proteins in either intrabody-TurboID condition (Permutation-based t-test FDR=0.01, s_0_=0.1). H. Top 3 GO:CC, MF or BP terms enriched in either synaptic condition. Numbers in parenthesis indicate the amounts of annotated proteins in each term. I. Correlation between the normalized physical distance of the plotted proteins between PSD95 and Homer1 (x-axis) and the normalized log2FC between PSD95-TurboID and Homer1-TurboID (y-axis). Dashed line is a fitted linear model based on proteins in red while the proteins in black are inferred based on this model. J to M. Differential enrichment of proteins with the mentioned GO terms in the Homer1-TurboID or PSD95-TurboID conditions. Abbreviations: CC: Cellular compartment, MF: Molecular Function, BP: Biological Process.

We used a previously optimized sample preparation workflow to pull down, elute and digest labeled postsynaptic proteins (Figure S2A; (Marcassa *et al*., 2025), see Methods). Data-independent acquisition (DIA), followed by DIA-NN analysis (Demichev *et al*., 2020) for label-free quantification (LFQ) identified 8320 proteins across all samples (Figure S2B). LFQ intensity distributions were comparable across Ctrl- and intrabody-TurboID samples (Figure S2C), as DIA-NN performs retention time-dependent cross-run normalization. Filtering background proteins, common contaminants and endogenously biotinylated proteins from the analysis resulted in 5128 proteins enriched in Ctrl-, PSD95- and Homer1-TurboID (Figure S2D). Principal Component Analysis (PCA) separated replicates according to experimental condition (Figure 2C). Comparing PSD95- and Homer1-TurboID conditions against Ctrl-TurboID yielded 1708 differentially enriched proteins (DEPs) at postsynapses (Figures 2D, S2D). Labeling by PSD95- and Homer1-TurboID resulted in distinct, but partially overlapping enrichment of synaptic proteins, with 961 and 1241 synaptic DEPs, respectively, of which 494 overlapped (Figure 2D).

Gene Ontology (GO) analysis (Subramanian *et al*., 2005; Yu *et al*., 2012; Liberzon *et al*., 2015) identified the Cellular Compartment (CC) terms “Postsynaptic Specialization” and “Neuron to Neuron Synapse” as most enriched in the PSD95- and Homer1-TurboID datasets, while the Ctrl-TurboID dataset was enriched in nuclear terms (Figure 2E). Of the DEPs for PSD95- and Homer1-TurboID, 33% and 23%, respectively, were annotated in SynGO, a synapse-centered GO analysis tool (Koopmans *et al*., 2019), and skewed towards the postsynaptic compartment (Figures 2F and S2E). Together, these results confirm the specificity of our approach and the robustness of our filtering pipeline.

Direct comparison of the 1708 postsynaptic proteins between PSD95- and Homer1-TurboID identified a total of 1450 DEPs (Figure 2G and S2D), with 915 proteins enriched in Homer1-TurboID and 535 proteins enriched in PSD95-TurboID. Top GO terms for CC, Molecular Function (MF) and Biological Process (BP) were remarkably dissimilar between the probes (Figure 2H): while PSD95-TurboID enriched for more cleft-proximal terms (e.g. “GO:CC Postsynaptic Specialization Membrane” and “GO:CC Ionotropic Glutamate Receptor Complex”), Homer1-TurboID enriched for more cleft-distal and signaling terms (e.g. “GO:MF GTPase Activator Activity” and “GO:BP Regulation of Organelle Assembly”).

These differences mirror structural observations showing that PSD95 and Homer1 occupy distinct subdomains of the PSD, with PSD95 positioned ∼40 nm and Homer1 ∼80 nm away from the synaptic cleft (Valtschanoff and Weinberg, 2001; Dani *et al*., 2010). Their respective intrabody-TurboID fusions would therefore be expected to preferentially label different subsets of synaptic proteins: cleft-proximal for PSD95-TurboID and cleft-distal for Homer1-TurboID (Figure S1A). To examine subsynaptic enrichment of proteins labeled by intrabody-TurboID fusions, we compared our MS results with the super-resolution microscopy dataset from Dani et al., which maps the relative position of postsynaptic proteins Dlg4 (PSD95), Grin2b, Gria1, Shanks, Camk2a and Homer1 at the synapse (Dani *et al*., 2010). The log_2_ fold change (log_2_FC) between PSD95-TurboID and Homer1-TurboID for each of these proteins strongly correlated with their normalized physical distances to PSD95 and Homer1 (R^2^=0.96; Figure 2I), indicating that the differential enrichment of proteins in our dataset reflects their relative physical localization within the PSD.

Consequently, inferred physical distances for other proteins in our dataset should predict their localization within the PSD. Indeed, we observed that the inferred position of proteins such as Dlgaps and Syngap1 aligns with the laminar organization of the PSD, in which they act as molecular bridges between Dlgs and Shanks (Valtschanoff and Weinberg, 2001; Sheng and Kim, 2011; Liu *et al*., 2019). Using this approach, we further predicted membrane localization for PSD-associated synaptic adhesion molecules in our dataset, including Lrrtm1, Nlgn1 and Lrrc4 (Figure 2I). Together, these results demonstrate a close correlation between differential protein enrichment in our intrabody-TurboID dataset and physical location within the PSD as determined by super-resolution microscopy.

GO analysis further supports the spatial specificity of the PSD95- and Homer1-TurboID probes. Proteins with cytoskeleton- and signaling-related GO terms were overly represented in the Homer1-TurboID condition (Figures 2J, 2K and S2E-I), while proteins related to glutamate receptors and plasma membrane were enriched in the PSD95-TurboID condition (Figures 2L, 2M and S2E-I), consistent with the cleft-distal to cleft-proximal axis we observe in our intrabody-TurboID dataset.

Most identified proteins are postsynaptic (Figures 2E, 2F and S2E). Despite stringent washes and filtering steps, we detected a small number of presynaptic proteins in the list of synaptic DEPs, including Slc17a7 (Vglut1), Sv2b, and Ptprs (Figure S2G). These proteins were predominantly present in the PSD95-TurboID condition, which is likely due to co-isolation of macromolecular complexes, including trans-synaptic partners, using this approach.

Taken together, PSD95-TurboID and Homer1-TurboID each label overlapping but distinct compartments of the PSD, mapping protein composition of CA1 PN excitatory inputs with deep proteome coverage and subsynaptic specificity.

### Intrabody-TurboID reveals extensive molecular heterogeneity of CA1 excitatory inputs

CA1 PNs integrate two major streams of information that are segregated along a proximal-distal dendritic axis. SCs from CA3 PNs target the proximal dendritic domains of CA1 neurons in the SO and SR laminae, whereas the TA pathway from layer III entorhinal cortex neurons targets the distal domain in SLM (Remondes and Schuman, 2002, 2004; Lisman and Grace, 2005; Kesner and Rolls, 2015; Valero and de la Prida, 2018).

Proximal and distal CA1 PN inputs differ in their composition of glutamate receptors and ion channels, which plays an important role in how CA1 PNs integrate synaptic inputs (Spruston, 2008). For example, distal inputs have a higher NMDAR/AMPAR ratio compared to proximal inputs (Otmakhova, Otmakhov and Lisman, 2002; Nicholson *et al*., 2006), compensating for their greater electrotonic distance from the soma. Distal inputs are also enriched in the hyperpolarization-activated cation channel Hcn1 (Lörincz *et al*., 2002; Notomi and Shigemoto, 2004), which shapes synaptic integration, dendritic excitability and the propagation of entorhinal cortex input (Magee, 1998).

To map the molecular architecture of CA1 PN inputs, we used the PSD95-TurboID probe for profiling of cleft-proximal synaptic proteins with laminar resolution. We injected PSD95- and Ctrl-TurboID AAVs in CA1 and micro-dissected the three synaptic laminae in dorsal CA1 in the PSD95-TurboID condition (Figure 3B). We verified construct expression and biotinylation levels by Western blotting (Figure S3A), and verified dissection accuracy by evaluating the levels of Nptx1, a synaptic protein previously shown to be enriched in proximal SO and SR relative to distal SLM (Figure S3A) (Cummings *et al*., 2017; Apóstolo *et al*., 2020).

**Figure 3.**
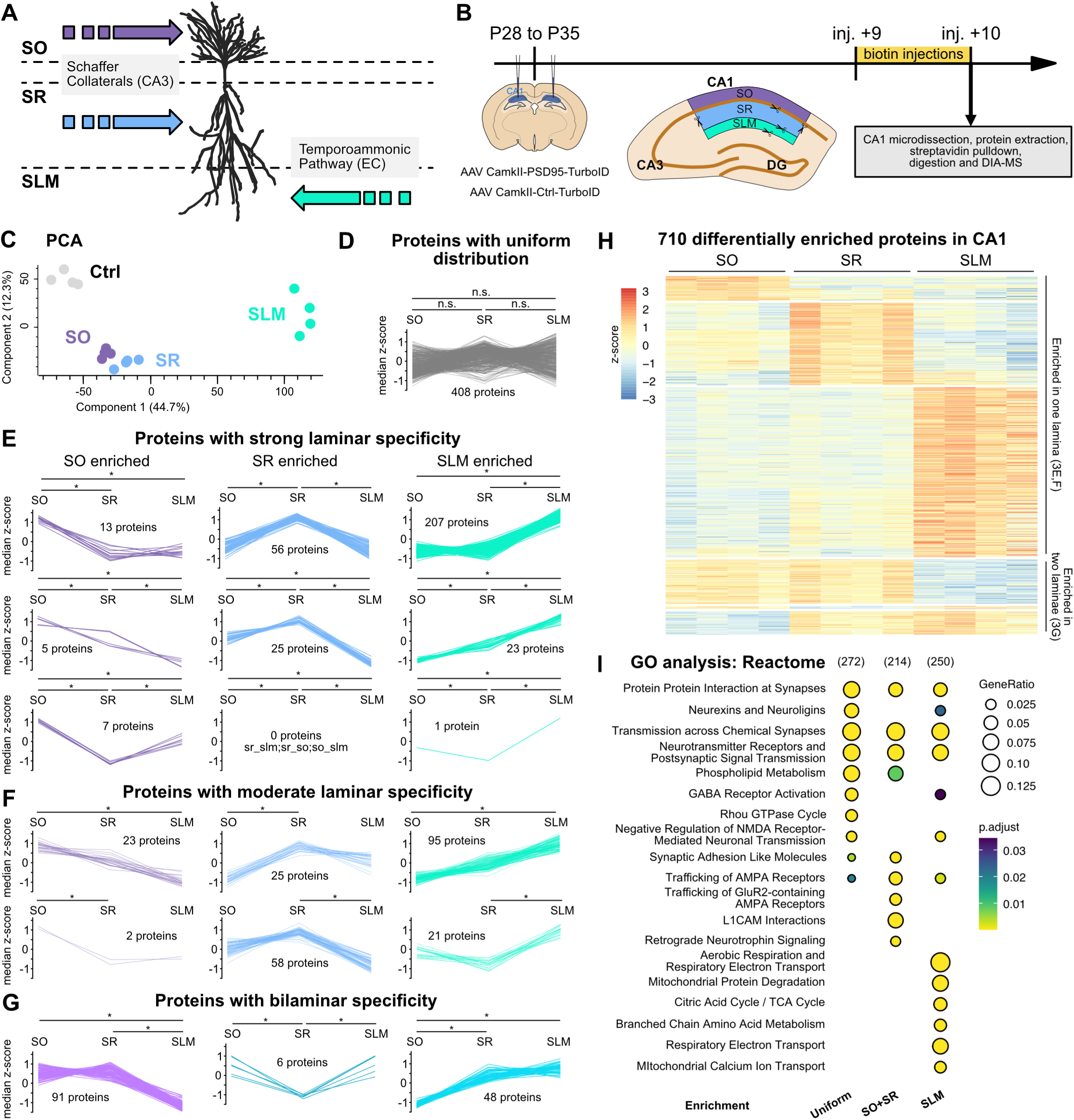
Lamina-specific synaptic composition in hippocampal CA1 pyramidal neurons. A. Schematic of the segregation of the main synaptic inputs onto CA1 pyramidal neurons. B. Experimental workflow (n=4 replicates per condition, 8 animals per replicate for PSD95-TurboID, 4 animals per replicate for Ctrl-TurboID, a mix of males and females). C. PCA showing separation of different replicates based on MS intensity. D. Distribution profile of non-significant proteins across the three CA1 laminae (ANOVA with Benjamini-Hochberg FDR=0.05 followed by Tukey’s HSD). E to G. Distribution profile of proteins strongly enriched in one lamina (E), moderately enriched in one lamina (F), or predominant in two laminae (G). H. Heatmap showing all 710 DEPs corresponding to the profiles shown in E-G. I. GO analysis of Uniform, SO+SR, and SLM enriched proteomes using Reactome terms. Numbers in parenthesis indicate the amounts of GO-annotated proteins in each group. Abbreviations: SO: *stratum oriens*, SR: *stratum radiatum*, SLM: *stratum lacunosum-moleculare*.

We identified a total of 8388 proteins (Figure S3B), with comparable intensities across TurboID samples (Figure S3C). PCA analysis revealed clear separation by lamina, with SO and SR clustering more closely, likely due to shared SC input from CA3 (Figure 3C). Our filtering pipeline yielded 1510 synaptic DEPs in the PSD95-TurboID condition relative to Ctrl-TurboID (Figure S3D-E). To exclude non-cell-autonomous proteins, we cross-referenced this list with single-cell transcriptomic data from the Allen Brain Cell (ABC) Atlas (Yao *et al*., 2023), retaining only CA1 PN-expressed proteins and excluding extracellular matrix proteins (Figure S3D, see Supplementary tables). This resulted in a final list of 1118 postsynaptic proteins across all laminae, 408 of which displayed a uniform distribution (Figure 3D and S3D). The other 710 proteins showed laminar enrichment (Figures 3E-H). Among these, we defined proteins enriched in one lamina relative to both others as having strong laminar specificity (Figure 3E), and proteins enriched in one lamina relative to one other lamina as displaying moderate laminar specificity (Figure 3F). Several proteins showed bilaminar specificity, with enrichment in two laminae relative to the third (Figure 3G).

To facilitate biological interpretation, we grouped proteins with strong or moderate laminar specificity into lamina-specific categories (SO, SR and SLM), and bilaminar proteins into SO/SR, SO/SLM and SR/SLM groups. These six groups capture the principal modes of laminar organization observed in our dataset (Figure 3H). The largest group was the SLM group (347 proteins), followed by SR (168 proteins) and SO (48 proteins). The bilaminar groups SO/SR, SR/SLM, and SO/SLM comprised 91, 48 and 6 proteins, respectively. The relatively high number of DEPs in SLM may reflect input diversity, as dorsal CA1 PN distal dendrites also receive a small proportion of thalamic inputs (Griffin, 2015), although this is debated (Andrianova *et al*., 2025). Alternatively, grouping by major presynaptic input shows that numbers of DEPs are similar between proximal SC inputs (SO+SR; 307 proteins) and distal TA inputs (SLM; 347 proteins).

To place these results in a biological context, we performed GO term analysis on the uniform group, the proximal group, and the distal group (Figures 3I and S3F, see Material and Methods). Top Reactome (Milacic *et al*., 2024) terms highlighted synaptic pathways across all groups (Figure 3I), while other terms showed domain-specific enrichment, such as “L1CAM interactions” and “Retrograde Neurotrophin Signaling” in the proximal group. Notably, terms related to mitochondrial function such as “Aerobic Respiration and Respiratory Electron Transport” were specifically enriched in the distal group. These observations were similar when performing GO analysis using CC and BP terms (Figure S3F). Thus, GO analysis reveals shared synaptic functions and domain-specific signatures, most prominently a pronounced enrichment of mitochondrial pathways in distal inputs.

Reactome GO analysis also identified “GABA Receptor Activation” in the uniform and distal groups (Figure 3I). The presence of inhibitory GABA_A_ receptors (GABAARs) is consistent with the presence of dually innervated CA1 spines (Iascone *et al*., 2020; Kleinjan *et al*., 2023); the proximity of SLM shaft synapses to inhibitory PSDs (Harris, Jensen and Tsao, 1992; Megías *et al*., 2001; Reilly, Hanson and Phillips, 2011), and activity-dependent trafficking of GABAARs to excitatory synapses (Luca *et al*., 2017).

Together, these results show that CA1 PN excitatory inputs are composed of distinct combinations of proteins. Having established the laminar organization of the CA1 PN postsynaptic proteome, we next asked how these molecular patterns relate to known functional differences between proximal and distal inputs.

### The molecular logic of CA1 PN excitatory inputs follows a proximal-distal axis

To understand how the molecular composition of CA1 PN excitatory inputs relates to their functional differences, we examined the laminar distribution of major classes of synaptic proteins, including glutamate receptors, ion channels, intracellular scaffolds, adhesion molecules, and regulators of protein dynamics, along the proximal-distal dendritic axis.

We first focused on glutamate receptors and their auxiliary subunits. AMPARs (Gria1-4) show strong SO or SR enrichment, as do most of their auxiliary proteins, including Shisa6 and the TARP (transmembrane AMPA receptor regulatory protein) family member Cacng3 (Figures 4A and S3G). In contrast, the NMDAR subunit Grin1 is enriched in SLM, consistent with known proximal-distal distributions of AMPARs and NMDARs in CA1 PNs (Otmakhova, Otmakhov and Lisman, 2002; Nicholson *et al*., 2006). Kainate receptor subunits (Grik2, Grik5) and their auxiliary proteins Neto1 and Neto2, are strongly biased toward SLM (Figures 4A and S3G). Finally, the delta glutamate receptors Grid1 and Grid2, which do not bind glutamate, are enriched in SLM. Together, these patterns reveal that proximal and distal inputs are defined by a specific complement of glutamate receptors and auxiliary proteins.

**Figure 4.**
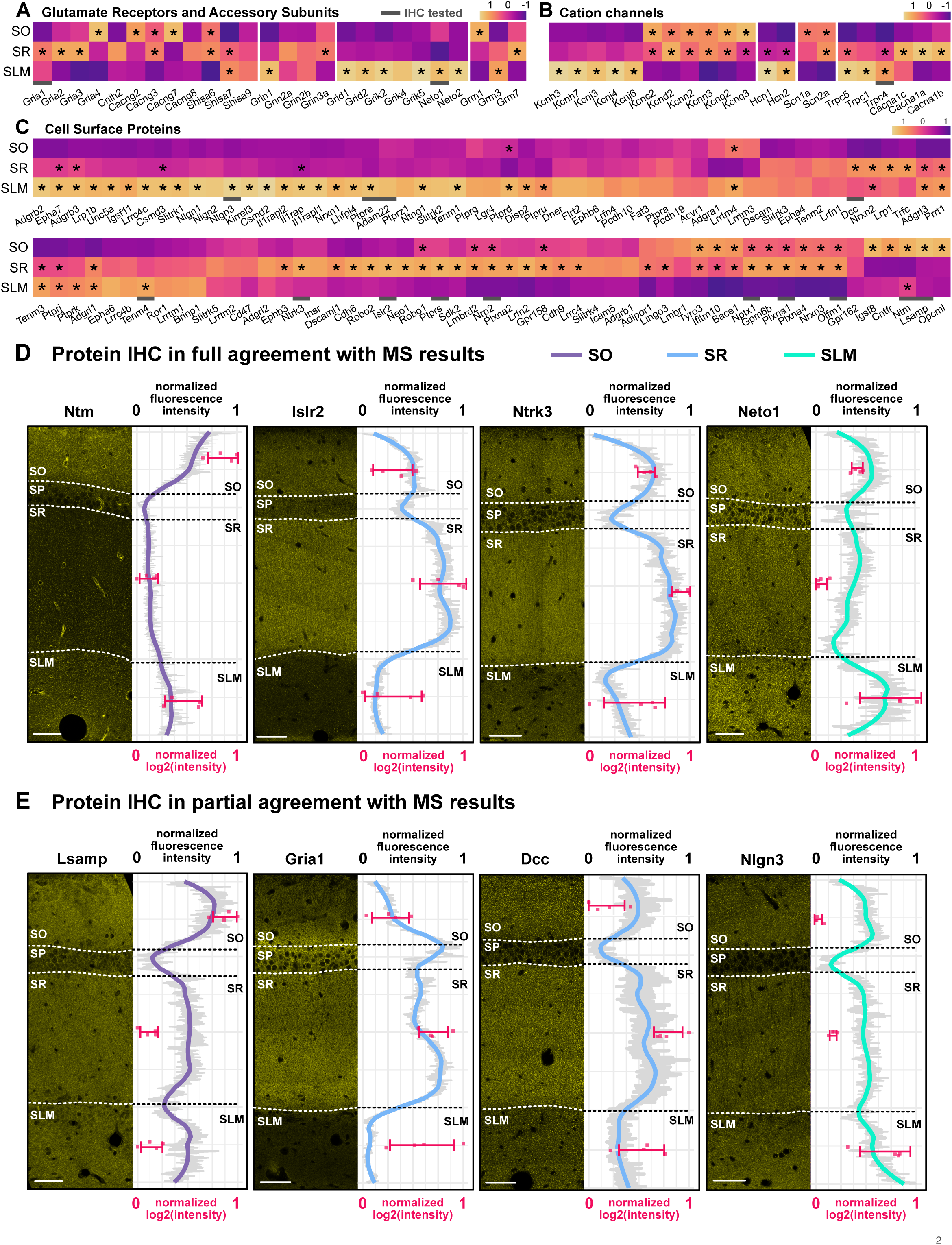
Validation of DEPs hippocampal CA1. A-C. Heatmaps showing the relative distribution across CA1 synaptic laminae of glutamate receptors (A), ion channels (B) or CSPs (C). D-E. Hippocampal CA1 stainings against different proteins in the laminar dataset. For each tested antibody, we show a representative confocal image (fire scale, single plane, left panels), and a plot of the normalized fluorescence intensity along the radial axis overlaid on a plot showing the normalized MS log_2_(intensity) (right panels). Abbreviations: SO: *stratum oriens*, SR: *stratum radiatum*, SLM: *stratum lacunosum-moleculare*.

Ion channels exhibit similarly striking differential enrichment along the proximal-distal dendritic axis (Figures 4B and S3G). As expected, cation channels Hcn1 and Hcn2 display a proximal-distal gradient (Lörincz *et al*., 2002; Notomi and Shigemoto, 2004). Potassium channel families segregate sharply: some are enriched in SO and SR (e.g. Kcnd2 (Kv4.2), Kcnn2 (SK2) and Kcnq2 (Kv7.2)), whereas others preferentially localize to SLM (e.g. Kcnh3 (Kv12.2), Kcnh7 (Kv11.3), Kcnj3 (Kir3.1) and Kcnj4 (Kir2.3). Calcium channels are also differentially distributed, with Cacna1c enriched in SR and Trp channels (Trpc1, 4 and 5) enriched in SLM (Figure 4B). These ion channel distributions underscore a domain-specific organization of excitability mechanisms across CA1 dendrites.

The differential distribution of proteins at segregated inputs likely arises from coordinated cell-autonomous and cell-non-autonomous processes. Intracellular and trans-synaptic scaffolds can anchor proteins to specific subcellular compartments (Meier *et al*., 2001; Nishimura-Akiyoshi *et al*., 2007; Uemura *et al*., 2010; Verpelli *et al*., 2012; Chen *et al*., 2015; Apóstolo and de Wit, 2019). To gain insight into the molecular basis of glutamate receptor and ion channel localization along the proximal-distal dendritic axis, we first examined intracellular scaffold protein distribution. Among core PSD scaffolds, few show strong laminar differences (Figure S4A). Camk2 family members are an exception, with several enriched in SLM. Additionally, we find Dlg5 and Dlgap4 enriched in SR, while Dlg1 (SAP97) and Dlg4 (PSD95) are enriched in SLM. Because our dataset is cross-run normalized, these differences likely reflect differences in average synapse composition rather than synapse density. For Dlg4 (PSD95), this aligns with prior findings that although SLM contains fewer excitatory synapses, dendritic spines and PSDs are larger, and PSD95 nanocluster intensity is higher (Megías *et al*., 2001; Broadhead *et al*., 2016; Zhu *et al*., 2018; Santuy *et al*., 2020). These results suggest that intracellular scaffolds contribute modestly to input-specific protein distributions.

We next examined trans-synaptic scaffolds (Feng and Zhang, 2009; Südhof, 2018), which include synaptic adhesion molecules and secreted scaffolding proteins collectively referred to as cell-surface proteins (CSPs). These molecules form trans-synaptic complexes that can specify neurotransmitter receptor composition (Pinan-Lucarré *et al*., 2014; Jang, Lee and Kim, 2017; Dai *et al*., 2022; Merrion *et al*., 2025). In contrast to intracellular scaffolds, CSPs show marked input-specific enrichment (Figure 4C). Leucine-rich repeat (LRR) receptors, immunoglobulin superfamily (IgSF) members, adhesion G-protein coupled receptors (adhesion GPCRs), neuroligin, plexin, ephrin and cadherin receptor families are all represented and display distinct laminar patterns along the proximal-distal dendritic axis (Figure 4C). Interestingly, although SO and SR both contain SC inputs from CA3 neurons, their CSP composition differs substantially. The IgLON (immunoglobulin Lsamp/Obcam/Ntm) family is particularly enriched in SO, with all identified members enriched in SO (Lsamp, Ntm and Opcml) (Figure 4C). Together, the distinct CSP repertoire of each lamina suggests that trans-synaptic interactions play a key role in defining input-specific properties.

Finally, intracellular protein dynamics, including local translation and protein degradation, can shape protein distribution within compartmentalized neurons (Holt, Martin and Schuman, 2019; Grochowska *et al*., 2022; Karpova *et al*., 2025). We find extensive differential enrichment of RNA-binding proteins across laminae (Figure S4C). In contrast, laminar differences in E3 Ubiquitin ligases are modest (Figure S4D). These findings point to local protein synthesis, rather than proteasomal degradation, as a dominant mechanism in shaping input-specific proteomes.

Together, these results show that the molecular signatures of CA1 PN excitatory inputs are defined by differential enrichment of glutamate receptors, ion channels, and CSPs. These distributions follow a proximal-distal logic and provide a molecular basis for the distinct functional properties of CA1 excitatory inputs.

### Validation of input-specific synaptic proteomes of CA1 PNs

To validate our laminar proteomic dataset, we performed immunohistochemistry (IHC) on mouse hippocampal CA1 sections using a panel of well-characterized antibodies against 20 proteins in the dataset (Figures 4D, 4E and S4E, Table S2). We compared the intensity profiles of antibody staining (from SO to SLM) with normalized LFQ intensities across CA1 laminae. The distribution of LFQ values for the SO-enriched protein Ntm closely matched its IHC intensity profile across CA1 (Figure 4D). Similarly, SR-enriched Islr2 and Ntrk3 and SLM-enriched Neto1 showed close agreement between laminar LFQ and IHC intensity profiles.

For some proteins, LFQ and IHC profiles showed partial divergence in one lamina but a similar enrichment trend, for example SR-enriched Gria1 and SLM-enriched Nlgn3 (Figure 4E). Based on these comparisons, we categorized the 20 analyzed proteins as showing full agreement (all three laminae match), partial agreement (two laminae or the enrichment trend match), or no agreement between proteomic and IHC results. Out of 20 proteins analyzed, 13 were in full agreement (Ntm, Neto1, Islr2, Ntrk3, Adam22, Flrt2, Olfm1, Ptprs, PlxnA1, Nptx1, Nrp2, Ptpre, Tenm4), 6 in partial agreement (Lsamp, Nlgn3, Gria1, Nos1, Trpc4, Dcc), and 1 did not agree (Cnr1, which is strongly enriched in axons in CA1 *stratum pyramidale* (SP), Figures 4D, 4E and S4E-F). Together, these results demonstrate strong correspondence between proteomic and immunohistological laminar profiles.

We further cross-referenced our findings with published studies employing orthogonal methods such as IHC and immunogold labeling (Table S1, Figure S4F). For example, Lrrc4 (NGL-2) and Lrrc4c (NGL-1) are established laminar markers for SR and SLM, respectively (Nishimura-Akiyoshi *et al*., 2007; Cummings *et al*., 2017), and their distribution in our dataset closely matched the reported patterns (Figure 4C). Using the same qualitative classification, we find that out of 43 proteins assessed in these studies, 22 are in full agreement, 8 are in partial agreement, 9 disagree and 4 are uncertain (e.g. due to limited image quality; Figure S4F) (Bekele-Arcuri *et al*., 1996; Lüscher *et al*., 1999; Shibata *et al*., 2003; Nicholson *et al*., 2006; Nishimura-Akiyoshi *et al*., 2007; Akashi *et al*., 2009; Linhoff *et al*., 2009; Tzingounis *et al*., 2010; de Wit *et al*., 2013; Klaassen *et al*., 2016; Cummings *et al*., 2017; Schroeder *et al*., 2018; Gutierrez, Dwyer and Franco, 2019; Sando, Jiang and Südhof, 2019; Apóstolo *et al*., 2020; Park *et al*., 2020; Uchigashima *et al*., 2020; Blockus *et al*., 2021; Matúš *et al*., 2024; Miao *et al*., 2024; Pelz *et al*., 2024; Meyer *et al*., 2025). Overall, the majority of proteins analyzed this way showed agreement between proteomic and histological data, further supporting the laminar signatures we identify.

In summary, these validations confirm the accuracy of our input-specific proteomic dataset, establishing it as a robust resource for linking molecular signatures to synaptic physiology.

### Proteomic profiling of CA1 inhibitory postsynaptic proteomes using Gphn-TurboID

In addition to receiving diverse excitatory inputs, CA1 PNs are innervated by multiple classes of inhibitory neurons, forming complex patterns of input (Tzilivaki *et al*., 2023). To characterize the molecular composition of inhibitory postsynapses, we extended the intrabody-TurboID approach using a FNIII-based monobody against the inhibitory scaffold gephyrin (Gphn) (Gross *et al*., 2013) (Figure 5A). In primary neuronal cultures, the Gphn-TurboID probe selectively labeled inhibitory postsynaptic proteins in the presence of biotin, as demonstrated by the colocalization of biotinylated proteins with Gphn (Figure 5B).

**Figure 5.**
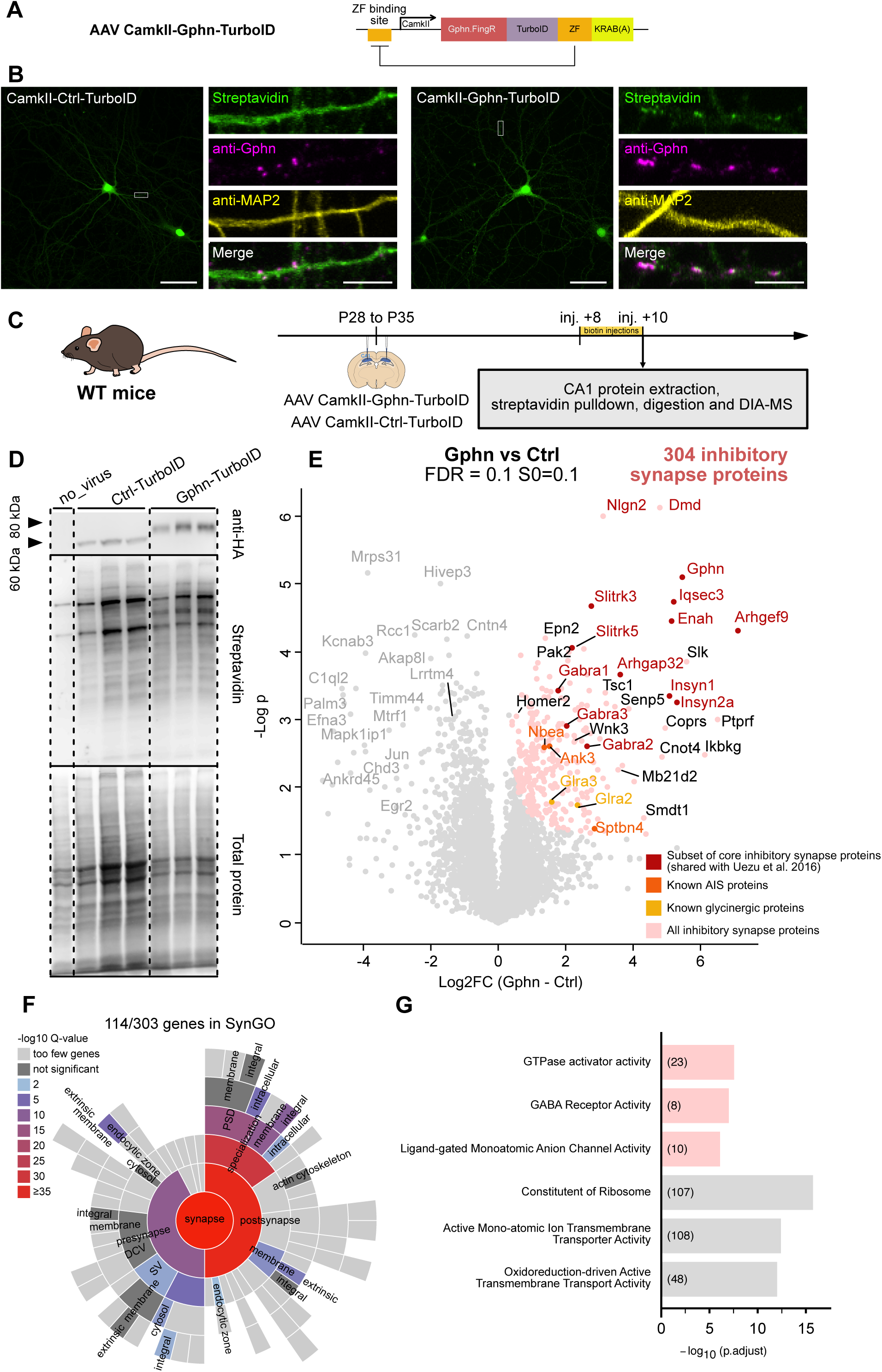
Gphn-TurboID efficiently biotinylates and identifies inhibitory synaptic proteomes. A. Schematic of the Gphn-TurboID AAV construct. B. Validation in primary neurons shows biotinylation localized to inhibitory synapses, as evidenced by low- and high-magnification z-stacks stained by streptavidin and an anti-Gphn antibody. C. Experimental workflow (n=3 replicates per condition, 3 animals per replicate, a mix of males and females). D. Western Blot showing expression of Gphn-TurboID and Ctrl-TurboID probes, as well as biotinylation levels for the three replicates in the experiment. E. Volcano plot showing the enrichment of inhibitory synaptic proteins in the Gphn-TurboID condition compared to control (Permutation-based t-test FDR=0.1, s_0_=0.1). F. SynGO enrichment diagram of the Gphn-TurboID enriched proteome. G. Top 3 GO:BP terms enriched in Gphn-TurboID and filtered proteomes. Numbers in parenthesis indicate the amounts of annotated proteins in each term. Scale bars are 50 μm for the low magnification images and 5 μm for the high-magnification images.

We next employed an experimental workflow analogous to that used for excitatory synapses, injecting CA1 with Gphn-TurboID and Ctrl-TurboID AAVs (Figure 5C). Western blotting confirmed appropriate construct expression and efficient biotinylation (Figure 5D). Proteomic analysis identified 6707 proteins across all replicates (Figure S5A), with comparable intensity distributions (Figure S5B) and clear PCA-based separation between Gphn-TurboID and Ctrl-TurboID conditions (Figure S5C). After filtering, we identified 304 inhibitory synapse proteins (Figures 5E and S5D). These included canonical inhibitory postsynaptic components, such as the scaffolds Gphn, Arhgef9 (Collybistin), and Insyn1/2, as well as the GABAAR subunits Gabra1/2/3 (Petrini *et al*., 2014; Tyagarajan and Fritschy, 2014; Uezu *et al*., 2016). We also identified inhibitory synapse-specific CSPs such as Nlgn2 and Slitrk3 (Varoqueaux, Jamain and Brose, 2004; Li *et al*., 2017), as well as glycinergic receptors Glra2/3 and axon initial segment (AIS)-associated proteins Nbea, Sptnb4 and Ank3 (Kubota *et al*., 2016; Hamdan *et al*., 2020). GO analysis further confirmed strong enrichment of postsynaptic terms (Figures 5F-G). Together, these results show that Gphn-TurboID effectively captures the molecular composition of inhibitory postsynapses in CA1 PNs.

A previous study used overexpression of a biotin ligase fused to several inhibitory baits, including Gphn and collybistin, to map inhibitory synapse composition in cortex and hippocampus (Uezu *et al*., 2016). Comparison of the two datasets revealed 39 shared proteins, which we designate as “core” inhibitory postsynapse components (Figure 5E, S5E). Notably, 265 proteins were only identified in our dataset, whereas 143 were unique to the Uezu et al. dataset, suggesting extensive inhibitory synapse diversity dependent on brain region, cell-type, or methodological differences (Kubota *et al*., 2016).

Together, these findings highlight both the molecular complexity of inhibitory postsynapses in CA1 PNs, and the modularity and broad applicability of the intrabody-TurboID approach across distinct synaptic compartments.

### Intrabody-TurboID reveals early and late phases of synaptic remodeling in a mouse model of AD

CA1 PNs are among the first neuronal populations to be affected in AD, raising the possibility that their inputs are particularly vulnerable (West *et al*., 1994; Furcila *et al*., 2019). In *App^NL-G-F^* mice, which express a humanized Aβ sequence with three familial AD mutations from the endogenous *App* locus (Saito *et al*., 2014), CA1 PNs begin to show defects in spontaneous excitatory synaptic transmission at 2 months of age (Calafate *et al*., 2023). These defects precede Aβ plaque deposition, apparent at 3 months in CA1 laminae (Rice *et al*., 2020). Synaptic transmission in CA1 PNs transiently normalizes due to homeostatic mechanisms but ultimately fails by 6 months, a stage characterized by extensive Aβ plaque deposition, neuroinflammation, and memory deficits (Saito *et al*., 2014; Rice *et al*., 2020; Calafate *et al*., 2023).

Changes in the molecular architecture of CA1 PN inputs at early stages of AD have been difficult to resolve. Previous proteomic studies either performed bulk tissue analysis in mouse models or sampled from post-mortem material from symptomatic stages of the pathology (Savas *et al*., 2017; Hesse *et al*., 2019; Bai *et al*., 2020; Levites *et al*., 2024; Pichet Binette *et al*., 2024). To investigate early vulnerability of CA1 postsynaptic inputs to Aβ pathology, we assessed their protein composition in *App^NL-G-F^* mice at presymptomatic (1 month) and symptomatic (6 months) stages (Figure 6A-C). DIA-MS analysis identified 8629 proteins across all conditions (Figure S6A-B). Filtering yielded 1221 synaptic proteins (Figure S6C) and 129 DEPs between *App^NL-G-F^* and WT mice across both timepoints (Figures 6D and S6C).

**Figure 6.**
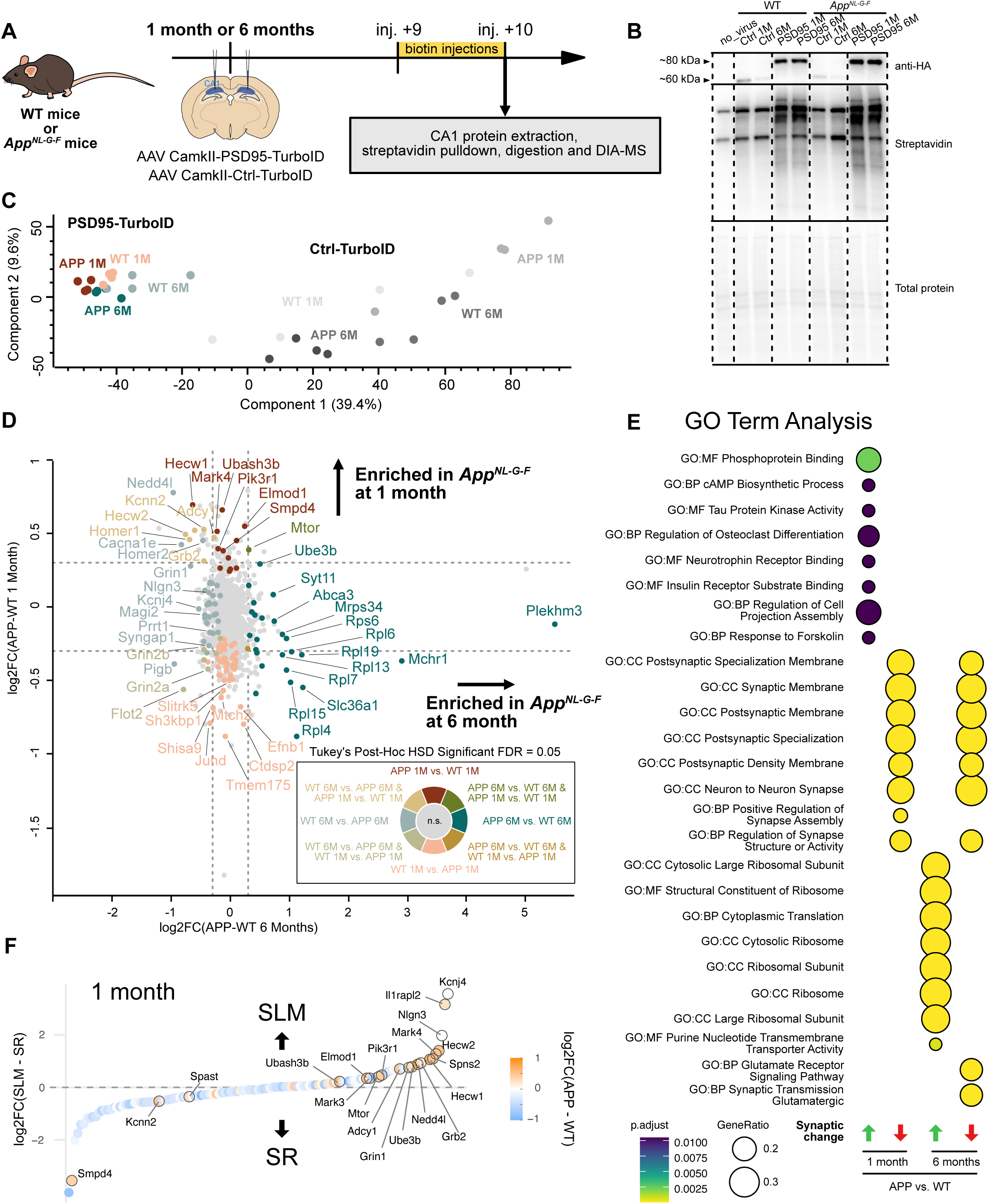
Intrabody TurboID quantifies early and late AD-related synaptic changes in a vulnerable neuronal population. A. Experimental workflow (n=4 replicates per condition, 3-4 animals per replicate, a mix of males and females). B. Western Blot showing expression of PSD95-TurboID and Ctrl-TurboID probes, as well as biotinylation levels for the two out of four replicates in the experiment. C. PCA showing the separation of the experimental conditions at 1 month (y-axis) and 6 months (x-axis) of age. Significant proteins are color-coded according to the legend (ANOVA with Benjamini-Hochberg FDR=0.05 followed by Tukey’s HSD). E. GO term analysis of synaptic DEPs at 1 month and 6 months between the AD and WT conditions. F. Rank plot comparing enrichment of AD synaptic proteins in SR and SLM.

At 1 month, *App^NL-G-F^* CA1 PNs already exhibited changes in synaptic protein composition. At this stage, 73 proteins were differentially enriched (19 upregulated, 54 downregulated). Upregulated proteins included signaling effectors (Mtor, Adcy1/2, Pik3r1, Grb2 and the tau kinases Mark3/4), E3 ubiquitin ligases (Hecw1/2), the scaffold Homer1 and the adhesion molecule Il1rapl2 (Figure 6D). Downregulated proteins included glutamate receptors and auxiliary subunits (Grin2a/2b and Shisa9) as well as CSPs (Lrrc4b, Efnb1, Dcc, Elfn2, Ntrk3, Adgrb3, Lrrtm1, Lrp1b and Slitrks). GO analysis mirrored these findings, showing enrichment of terms such as “GO:MF Phosphoprotein Binding”, “GO:BP cAMP Biosynthetic Process” and “GO:MF Tau Protein Kinase Activity” for upregulated proteins, and “GO:BP Positive Regulation of Synapse Assembly” for downregulated proteins (Figure 6E). Together, these changes suggest an early phase of synaptic remodeling and compensation in CA1 PNs in *App^NL-G-F^* mice at 1 month. Notably, these changes occur before detectable alterations in synaptic transmission (Calafate *et al*., 2023).

At 6 months, 70 proteins were differentially enriched (27 upregulated, 43 downregulated). Upregulated proteins included Mtor, regulators of endolysosomal and autophagy pathways (Clcn6, Clcn7, Syt11 and Atg9a), and multiple ribosomal proteins (Figure 6D). Downregulated proteins included glutamate receptors (Gria1, Grin1/2a/2b and Grm1), synaptic scaffolds and effectors (Syngap1, Prrt1, Magi2 and Homer1/2), CSPs (Nlgn3, Unc5a and Adam22), and signaling effectors (Rgs17, Dock4, Adcy1, Grb2). GO analysis showed upregulation of ribosome- and translation-related terms and downregulation of glutamatergic synaptic transmission-related terms (Figure 6E).

Remarkably few DEPs overlapped between the two timepoints, suggesting distinct early and late phases of the synaptic response to Aβ. Only 7 proteins were altered at both ages: Mtor (upregulated), and Grin2a/2b, Flot2, Begain, Slc7a14 and Cotl1 (downregulated) (Figure 6D). Several proteins that were upregulated at 1 month, such as Ubash3b, Mark3/4, Hecw1/2, Adcy1, Grb2, Kcnn2 and Homer1, were unchanged or downregulated at 6 months. The changes in synaptic proteins at 6 months suggest that the early phase of synaptic remodeling in *App^NL-G-F^* mice is not maintained; yet, systemic downregulation of multiple proteins related to glutamatergic transmission and signaling is still prevalent. These changes are parallelled by an upregulation of ribosomal proteins (Figure 6D), possibly reflecting compensatory proteostatic mechanisms at this stage.

Interestingly, by cross-referencing the log_2_FC of the 129 AD DEPs with LFQ values in our laminar dataset, we find that many of the upregulated synaptic proteins at 1 month are skewed towards SLM compared to SR (Figure 6F), potentially reflecting the heightened vulnerability of this input in AD (West *et al*., 1994).

Taken together, we identify distinct phases of CA1 PN postsynaptic remodeling in response to Aβ pathology in *App^NL-G-F^* mice: an early phase of remodeling and compensation that precedes changes in synaptic transmission, followed by a later phase marked by downregulation of glutamatergic transmission-related proteins and upregulated translation machinery.

## Discussion

Categorizing the molecular heterogeneity of synaptic connections at the cell type- and input type-level is fundamental to elucidate how neural circuits are established, maintained, and perturbed in disease (O’Rourke *et al*., 2012; Cizeron *et al*., 2020; van Oostrum and Schuman, 2025). Pyramidal neurons, with their stereotyped input segregation, represent the basic computational units of the vertebrate brain. Among those, CA1 PNs are central to learning and memory and are a vulnerable population in neurodevelopmental and neurodegenerative disorders (West *et al*., 1994; Clement *et al*., 2012; Furcila *et al*., 2019; Sawicka *et al*., 2019; Jones *et al*., 2024). Here, our goal was to create a minimally invasive, versatile and scalable method to resolve synaptic proteomes in defined neuronal populations such as CA1 PNs. We show that the intrabody-TurboID approach profiles the molecular composition of CA1 PN inputs with high resolution and can be adapted to different binder types and subcellular compartments. Using this approach, we uncover a proximal-distal molecular logic that underlies CA1 PN input-specific properties and capture progressive changes in synaptic composition in *App^NL-G-F^* mice. Our results provide a strong case study for the systematic, high-resolution mapping of synaptic protein composition and dynamics during development, plasticity, and disease.

By targeting two postsynaptic baits (PSD95 and Homer1) with distinct distributions in the PSD, our approach achieves subsynaptic resolution, enabling analysis of excitatory postsynaptic organization. Using these baits as anchor points and cross-referencing our proteomic data to a super-resolution mapping study, we recapitulate the laminar organization of the PSD (Valtschanoff and Weinberg, 2001; Dani *et al*., 2010). We demonstrate that we can infer relative protein localization within the PSD based on differential enrichment of proteins in our dataset. As expected, glutamate receptors and CSPs are predicted to be cleft-proximal, while cytoskeleton regulators and signaling molecules are predicted to be more distal, consistent with the described organization of the PSD (Valtschanoff and Weinberg, 2001; Dani *et al*., 2010; Liu *et al*., 2019). This approach can be readily applied to interrogate PSD architecture across specific inputs or experimental conditions (e.g. changes induced by synaptic plasticity), using DIA-MS analysis as a proxy for spatial proximity. Furthermore, it highlights the modular nature of the approach: different intrabody-TurboID fusions can be used independently or in combination, depending on the desired spatial resolution and proteomic coverage. The PSD95 and Gphn binders we use here have been thoroughly validated (Gross *et al*., 2013; Rimbault *et al*., 2024), but an in-depth characterization of the Homer1 nanobody beyond qualitative assessments has not yet been performed.

We use PSD95-TurboID to map the molecular composition of the major excitatory inputs onto CA1 PNs, identifying distinct molecular signatures for each input that are characterized by differential enrichment of glutamate receptor subunits, ion channels and CSPs. Our data shows that molecular composition of CA1 inputs is organized along the proximal-distal dendritic axis, with proximal (SO and SR) inputs exhibiting more similarity to each other than to distal (SLM) inputs. We identify proximal and distal input-specific compositions of glutamate receptors and ion channels that reflect functional properties of CA1 PN inputs along the dendritic axis (Dudman, Tsay and Siegelbaum, 2007; Piskorowski and Chevaleyre, 2012; Larkum *et al*., 2022).

In proximal SO and SR, the combined action of fast AMPAR currents (through enrichment in AMPARs), NaV-driven EPSP boosting (through Scn2a), activity-induced repolarizing A-type and M-type K⁺ currents (through members of Kcnd and Kcnq families, respectively), and calcium-activated SK-currents (through Kcnn2/3), supports brief, temporally precise and segregated responses, suggesting that the proximal input is the main driver of somatic output (Hoffman *et al*., 1997; Stackman *et al*., 2002; Ngo-Anh *et al*., 2005; Peters *et al*., 2005; Kim *et al*., 2007; Spratt *et al*., 2019; Baculis, Zhang and Chung, 2020).

Synaptic signals are strongly attenuated the further away from the soma they are generated. In contrast to proximal inputs, distal inputs are considered modulatory and follow different activation rules: in SLM inputs, strong shunting *I_h_* currents (through Hcn channels), slowly deactivating voltage-gated currents (through Kcnh channels), and inwardly-rectifying currents (through Kcnj channels) are expected to stabilize the resting membrane potential (Magee, 1998; Lörincz *et al*., 2002; Chen and Johnston, 2005; Tsay, Dudman and Siegelbaum, 2007; Luján, Maylie and Adelman, 2009; Zhang *et al*., 2010), and attenuate the amplitude of EPSPs. In parallel, enrichment of kainate receptors and Trpc cation channels is a likely mechanism leading to prolonged EPSPs with a sustained Ca^2+^ component (Castillo, Malenka and Nicoll, 1997; Frerking and Ohliger-Frerking, 2002; Kim *et al*., 2003; Lerma and Marques, 2013; Malik and Johnston, 2017). This combination results in events with relatively small-amplitudes and prolonged kinetics at distal inputs, leading to reinforced temporal summation and a wider integration window with proximal inputs, in line with their role in coincidence detection (Otmakhova, Otmakhov and Lisman, 2002; Bittner, Andrasfalvy and Magee, 2012).

The molecular signatures of CA1 PN inputs provide insight into the underlying organization of these proximal-distal distributions of glutamate receptors and ion channels. Differences between proximal and distal inputs were most apparent in their CSP composition, whereas intracellular scaffolds did not show major differences between inputs. These findings suggest that extracellular interactions play a key role in defining input-specific properties (Pinan-Lucarré *et al*., 2014; Jang, Lee and Kim, 2017; Dai *et al*., 2022; Merrion *et al*., 2025). In addition to proximal-distal differences, our approach also resolves molecular differences between the proximal (SO and SR) input proteomes, which primarily differ in their CSP composition. Given the key role of this class of proteins in specifying circuit connectivity, these differences could reflect the fact that SO receives more inputs from distal CA3 and CA2 compared to SR (Ishizuka, Weber and Amaral, 1990).

Our results also reveal an enrichment of mitochondrial proteins in distal CA1 PN inputs (Figures 3I, S3F and S4B). This is an unexpected observation, as mitochondria are seldom localized in dendritic spines (Sorra and Harris, 2000; Faitg *et al*., 2021). However, CA1 dendrites in SLM have a higher proportion of shaft synapses (Megías *et al*., 2001), which are more likely to be in close proximity to mitochondria. Additionally, these organelles are morphologically and functionally distinct between laminae: proximal mitochondria are smaller and more fragmented, whereas distal mitochondria are longer (Virga *et al*., 2024) and more likely to have a higher efficiency of their respiratory chain, explaining enrichment in TCA cycle proteins (Gomes, Benedetto and Scorrano, 2011). Given recent studies implicating mitochondria in intracellular Ca^2+^ signaling (Hirabayashi *et al*., 2017; O’Hare *et al*., 2022; Virga *et al*., 2024), the SLM input-specific enrichment of mitochondrial proteins might reflect a role in the functional properties of distal synaptic inputs.

Like CA1 PNs, cortical PNs function as signal integrators and display a similar proximal-distal input organization that enables processing of bottom-up and top-down signals on proximal and distal dendritic domains, respectively (Spruston, 2008; Larkum *et al*., 2009; Takahashi *et al*., 2016; Suzuki and Larkum, 2020). It will be of interest to determine whether common molecular signatures shape differential synaptic properties of proximal and distal inputs across cortical and hippocampal PN types.

Taken together, the CA1 PN input-specific synaptic proteomes we identify reveal how glutamate receptors, auxiliary subunits, ion channels, trans-synaptic interactions and signaling proteins specify functional differences along the proximal-distal dendritic axis (Figure S7). This dataset provides a foundation for building proteome-informed models of neuronal processing and signal integration in CA1 PNs (Nandi *et al*., 2022; Romani *et al*., 2024). Compared to a hippocampal FASS dataset (Kaulich *et al*., 2025), the advantage of our approach is the cell-autonomous aspect of the identified synaptic proteomes. Ultimately, these two techniques represent complementary strategies to map molecular signatures of synapses in a systematic manner across the synaptic interface. Here, we have used the excitatory CamkIIα(0.4) promoter which can exhibit leakiness (Veres *et al*., 2023), but given that CA1 is mostly composed of excitatory neurons (>85%, (Megías *et al*., 2001)), this effect is likely minimal. To restrict selectivity to defined PN subtypes, future studies could use Cre-dependent strategies.

Finally, we uncover early changes in CA1 PN excitatory input proteome composition in response to Aβ pathology in *App^NL-G-F^*mice. Most downregulated synaptic proteins at both presymptomatic (1 month) and symptomatic (6 months) stages were glutamate receptors, auxiliary subunits, and CSPs. In contrast, upregulated proteins showed major differences between the two timepoints: the presymptomatic stage is characterized by changes in proteins of major signaling pathways, such as Mtor and the MAPK pathway (through Grb2 involved in signal transduction downstream of receptor tyrosine kinases) and second messenger pathways (cAMP through adenylate cyclases or IP3 through Pik3r1), whereas the symptomatic stage displays a strong upregulation of ribosomal proteins. These changes suggest an early phase of signaling-driven synaptic remodeling and compensation, followed by a later phase of translational compensation in CA1 PN synapses in *App^NL-G-F^* mice.

It will be of interest to determine how these changes in synaptic protein composition are related to functional alterations in CA1 PNs in *App^NL-G-F^* mice during early disease stages. We previously found that defects in spontaneous synaptic transmission in these neurons start at 2 months (Calafate *et al*., 2023). The proteomic changes we detect at 1 month could suggest that CA1 synapses are employing compensatory mechanisms to maintain synaptic transmission. Alternatively, subtle synaptic defects that are not apparent when analyzing spontaneous transmission may already be present at this early stage. Cross-referencing of our *App^NL-G-F^* CA1 PN dataset with the input-specific dataset revealed that proximal and distal inputs could be differentially affected at 1 month, but functional changes in distal inputs may not be apparent as their influence on spontaneous transmission recorded at the soma is small. Among the upregulated proteins at 6 months, we detect Mchr1, the receptor for melanin-concentrating hormone (MCH), a neuropeptide produced in lateral hypothalamic neurons that project to CA1 and other brain regions (Bittencourt *et al*., 1992; Calafate *et al*., 2023). The MCH system becomes progressively impaired during early disease stages in *App^NL-G-F^* mice, with MCH mRNA upregulated in hypothalamic axons of 3.5 months old mice (Calafate *et al*., 2023). Interestingly, MCH peptide levels in the brain of another AD mouse model are decreased at 6 months (Levites *et al*., 2024). The upregulation of Mchr1 in CA1 PN synapses at the symptomatic stage could reflect a compensatory response to decreased MCH levels.

Taken together, we detect early and progressive changes in CA1 PN excitatory inputs in *App^NL-G-F^* mice. While these alterations may arise from direct effects of Aβ on the postsynaptic compartment, indirect effects are also likely, given that Aβ disrupts presynaptic function (Hark *et al*., 2021; Jeans *et al*., 2025) and inhibitory synapses (Latif-Hernandez *et al*., 2020; Li *et al*., 2021). Nevertheless, these findings demonstrate the sensitivity of the intrabody-TurboID approach for detecting subtle, progressive synaptic alterations, providing a foundation for the systematic mapping of synaptic vulnerability in neurodegenerative disease models.

In conclusion, our work establishes intrabody-guided TurboID proteomics as a versatile approach for linking synaptic protein architectures to specific input organization in both physiological and disease contexts.

## Material and Methods

### Animals

All animal experiments were conducted according to the KU Leuven ethical guidelines and approved by the KU Leuven Ethical Committee for Animal Experimentation. Mice were maintained under standard housing conditions with continuous access to food and water. All mice in this study were maintained on 14 h light, 10 h dark light cycle from 7 to 21 h. Wild-type (WT) C57BL/6 J (obtained from JAX) and *APP^NL-G-F^* mice used in this study were either 4–5-weeks old at the time of injection or approximately 6 months. Genotypes were regularly checked by PCR analysis. Animals were either euthanized through anesthesia with isoflurane followed by decapitation (for proteomics experiments), or by injection with an irreversible dose of ketamine– xylazine (for perfusions).

### Plasmids

TurboID sequence was a gift from Alice Ting (Addgene plasmid #107169). Xph20 monobody plasmid was a gift from Matthieu Sainlos (Addgene plasmid # 133024). HC87 nanobody sequence a gift from James Trimmer (Addgene plasmid #135223). GPHN-FingR plasmid was a gift from Don Arnold (Addgene plasmid #46296). TurboID was fused at the C-terminus of the intrabodies and before the zinc finger (ZF) and KRAB repressor domains in the pCAG-GPHN-FingR plasmid by Gibson assembly (NEB) to yield pAAV-CAG-intrabody-plasmids, or directly after the transcriptional start site to yield pAAV-Ctrl-TurboID. An HA-tag was added C-terminally. The CAG promoter was swapped with a short CamkII promoter, resulting in the pAAV-CamkII plasmids described. All plasmids were grown in NEB Stable bacteria at 31C. Plasmids will be deposited at www.addgene.org or made available upon request.

### Cell lines

HEK293T-17 human embryonic kidney cells (thereafter referred to as HEK cells) were obtained from American Type Culture Collection (ATCC) cat# CRL-11268. HEK293T-17 cells were grown in Dulbecco’s modified Eagle’s medium (DMEM; Thermo Fisher Scientific) supplemented with 10% fetal bovine serum (FBS; Thermo Fisher) and penicillin/streptomycin (Thermo Fisher).

### Primary neuronal cultures

Neurons were cultured from E17.5 to E18.5 WT C57BL/6J mice embryos (cortical and hippocampal). Briefly, dissected hippocampi and cortices were incubated with trypsin (0.25% v/v, Thermo Fisher) in HBSS supplemented with 10 mM HEPES (Thermo Fisher) for 15 mins at 37C. After trypsin incubation, cortices were washed with MEM supplemented with 10% (v/v) horse serum (Thermo Fisher) and 0.6% (wt/vol) glucose (MEM-horse serum medium) 3 times. The cells were mechanically dissociated by repeatedly pipetting the tissue up and down in a flame-polished Pasteur pipette and then plated on poly-D-lysine (Alamanda Polymers) and laminin (Sigma-Aldrich) coated glass coverslips in 12 well cell-culture plates containing MEM-horse serum medium. After 2 to 4 h, after neurons attached to the substrate, the coverslips were flipped onto an astroglial feeder layer in 12-well plates containing neuronal culture medium with the following composition: neurobasal medium supplemented with B27 (1:50 dilution), 12 mM glucose, Glutamax, penicillin/streptomycin (1:500 dilution), 25 μM β-mercaptoethanol (all reagents from Thermo Fisher), and 20 μg/mL insulin (Sigma). Neurons grew face down over the feeder layer but were kept separate from the glia by plastic feet extruded from the 12-well plate using flame-heated forceps. To prevent overgrowth of glia, neuron cultures were treated with 10 μM 5-Fluoro-2′-deoxyuridine (Sigma-Aldrich) after 3 d. Cultures were maintained in a humidified incubator of 5% CO_2_/95% (v/v) atmosphere at 37°C. For the biotinylation experiments, cortical neurons were transduced with low-titer AAVs (1:1000 dilution) at DIV7. At DIV16/17, biotin was applied to the wells to a final concentration of 500 uM for 1 h.

### Low-titer AAV purification

Low-titer AAV production (used in primary neuronal cultures) was performed from cell culture supernatant. Briefly, HEK cells were plated in 12 well cell culture and upon reaching ∼80% confluency were transfected with a DNA::PEI complex consisting of 800 ng pDelta F6 plasmid, 400 ng RepCap 2/1 plasmid and 400 ng AAV genome plasmid, as well as 5.7 μl PEI, all in a total volume of 100 μl Optimem (Thermo Fisher). 3 days later, the supernatant was collected, filtered through a 0.45 um filter and concentrated by centrifugation on 100 kDa MWCO 0.5 ml Amicon columns (Sigma) to a volume of approximately 30 μl. The resulting AAVs were used at a dilution of 1:500 to 1:2000 to transduce primary neuronal cultures.

### High-titer AAV purification

High titer AAV production and purification was carried out as previously reported (Challis *et al*., 2019). Briefly, 6 plates of 70% confluent HEK cells were transfected with 20 μg of pDelta F6 plasmid, 10 μg of RepCap 2/9 plasmid and 10 μg AAV genome plasmid per plate using PEI transfection. 3 days later cells were collected by scraping and centrifugation. AAV particles in the supernatant were precipitated by PEG and added to cell lysates. Lysates were treated by Benzonase nuclease to remove cellular DNA. Lysates were loaded onto a iodixanol gradient (60%, 40%, 25%, 15%) and ultracentrifuged 1:40 h at 50.000 RPM at 12 °C. The 40% fraction containing the purified AAV was carefully collected and desalted and concentrated on a 100 kDa MWCO 4 ml Amicon column (Sigma). AAV purity was tested by silver staining using the ProteoSilver Silver Stain Kit (Sigma PROTSIL1-1KT). AAV titer was determined by qPCR to normalize titers between different viral vector batches using LightCycler 480 SYBR Green I Master (Roche 04707516001).

### Immunofluorescence (neuronal cultures)

Cultured neurons on coverslips were briefly washed with ice-cold PBS and fixed for 5 min with 4% PFA in PBS, then washed again 2x with PBS. Next, they were permeabilized and blocked using using PBS with 10% normal horse serum (NHS) and 0.25% Triton-X 100 for 30 min. Primary antibody incubation was done by flipping coverslips on parafilm in a droplet of antibody mix in PBS with 2% NHS and 0.05% Triton-X 100. This incubation step was performed overnight at 4C. Then coverslips were washed 3x with PBS and incubated with secondary antibodies in the same buffer and fashion as primary antibodies, but at RT for 3h. Coverslips were once again washed 2x with PBS, after which they were incubated in a DAPI solution in PBS (1:10000 dilution), and finally washed once more in PBS. Coverslips were mounted on microscope slides with Mowiol-4-88 (Millipore). Imaging was performed on a Nikon AX confocal system. For visualization purposes, brightness and contrast was adjusted in FIJI (Schindelin *et al*., 2012).

### Immunohistochemistry (brain sections)

Mice were anesthetized by intraperitoneal injection with a lethal dose of anesthetics (1 μl/g xylazine (VMB Xyl-M 2%), 2 μl/g ketamine (Eurovet Nimatek, 100 mg/ml), and 3 μl/g 0.9% saline) and transcardially perfused with 4% PFA in PBS. Brains were dissected, postfixed in 4% PFA in PBS at 4 °C overnight, washed in PBS and sliced to 80 μm thickness by vibratome (Campden Instruments 7000 smz 2). Sections were blocked in 10% NHS, 0.5 M Glycine, 0.5% Triton-X 100, 0.02% gelatin in PBS for 2 h before incubation with antibodies in 5% NHS, 0.5% Triton-X 100, 0.02% gelatin in PBS overnight at 4C. After extensive washing with 0.05% Triton-X 100 in PBS, sections were incubated in a similar secondary antibodies solution for 3 h. After another round of extensive washing, sections were counterstained with DAPI before being mounted on microscope slides with Mowiol. Imaging was performed on a LSM900 confocal microscope with Airyscan (Zeiss). For visualization purposes, brightness and contrast was adjusted in FIJI. Intensity profiles in Fig.4 were measured on raw images by selecting a fixed width ROI encompassing the image and using the “Plot Profile” function in FIJI.

### Antibodies

All primary antibodies used for immunofluorescence, immunohistochemistry or WBs in this study are described in Table S2. Secondary antibodies are fluorophore- or HRP-conjugated from Thermo Fisher and Jackson Laboratories and used at a dilution of 1:750 (IF), 1:500 (IHC) or 1:5000 (WB).

### Stereotactic injections

P28 mice were anesthetized with 5% isoflurane. Duratear artificial tears was applied to the eyes to prevent them from drying out. Mice were placed in a mouse stereotact (KOPF) on a hot plate kept at 37 °C. During the rest of the procedure 2.5% isoflurane was constantly administered. After shaving and disinfecting the mouse’s head, local anesthesia was administered by a subcutaneous injection with 6mg/kg lidocaine (xylocain 1%). An incision was made on the skin to reveal the skull. A burr hole was drilled for hippocampal CA1 (at coordinates x: 1.7, y: 2.2 from bregma, z: 1.25 from the surface of the brain) and 300 nl of AAV mix injected using a Nanoject III (Drummond) through a beveled capillary at 5 nl/s. TurboID AAVs were also co-transduced with an AAV expressing a fluorescent protein (AAV-hSyn-GFP in this case) in order to identify the injection location and spread in each animal. After a 5 min recovery, the capillary was pulled out at ∼0.02 mm/s. The incision was stitched with surgical glue (Millpledge Veterinary). After 6 h, mice health was examined and the animals were injected with 0.1 mg/kg buprenorphine. One injection per hemisphere was performed for every experiment. All AAVs were injected at concentration between 10 ^ 11 and 10 ^ 12 GC/mL.

### Brain lysate preparation

Nine and ten days after AAV injection, mice were injected subcutaneously with 1 ml of 2 mg/mL biotin in 1X PBS, adjusting the pH to 7.4 to increase biotin solubility. 3 to 5 h after the last injection, mice were briefly anaesthetized by isoflurane and brains collected in ice cold HBSS + 10 mM HEPES. For experiments described in Figures 2, 5, and 6, 500 μm coronal brain sections were produced using an ice-cold brain matrix (AgnThos 69-2165-1), and CA1 was dissected under a fluorescence stereomicroscope using fluorescent signal expressed by infected cells and morphological landmarks. Typically, 2/3 animals were used per replicate for these experiments. For experiments in Figure 3, because of hippocampal curvature, we could not use brain matrix sections to reliably separate laminae along the dorso-ventral axis, so we opted for thinner and more reproducible vibratome sections of 300 um using a Leica VT1200S. Since cutting vibratome sections takes significantly longer than sectioning using the brain matrix, these experiments were performed in carboxygenated ACSF to preserve cell and tissue integrity (Bischofberger *et al*., 2006)(87 mM NaCl, 25 mM NaHCO3, 10 mM Glucose, 75 mM Sucrose, 2.5 mM KCl, 1.25 mM NaH2PO4, 0.5 mM CaCl2, 7 mM MgCl2, 1mM Kynurenic acid, 5 mM Ascorbic acid, 3 mM Pyruvic acid, equilibrated with 5%CO_2_/95%O_2_, osmolarity ∼325 mOsm). CA1 laminae were dissected under a fluorescence stereomicroscope using fluorescent signal and morphological landmarks. To further decrease dissection time, we pooled SO and the *stratum pyramidale* (SP), as the SP layer containing neuronal cell bodies is extremely thin compared to the synaptic laminae, but is virtually devoid of excitatory inputs (Valero *et al*., 2015; Broadhead *et al*., 2016). Typically, 6 to 8 animals (a mix of males and females) were used per replicate for the PSD95-TurboID constructs. For Ctrl-TurboID injected mice in this experiment, we collected whole CA1 in the same fashion.

Tissue pieces from each mouse were collected in a separate tube, flash-frozen in liquid nitrogen after dissection buffer removal, and stored at −80 °C until needed. For protein extraction and pulldown, we extensively optimized published protocols (Branon *et al*., 2018; Marcassa *et al*., 2025). Tissue from one mouse was homogenized and lysed in 50 μl 1% SDS RIPA buffer (50 mM Tris pH 8, 150 mM NaCl, 1% SDS, 0.5% Sodium Deoxycholate, 1% Triton-X 100, 1X protease inhibitors) using a small glass Dounce homogenizer. Lysates were sonicated in a bath sonicator (124-9721 RS-online) at max power three times for 1 min with 30 s rest on ice in between. Lysates were then boiled at 95 °C for 5 min to dissociate the PSD and trans-synaptic complexes and immediately diluted to 0.5% SDS with SDS-free RIPA buffer. This step leads to the release of trans-synaptic complexes and increases the specificity of the pulldown, as reported in (Loh *et al*., 2016). After 3 h incubation at 4 °C with rotation, lysates were cleared by centrifugation at 20.000 G for 15 min at 4 °C and supernatant collection. Bicinchoninic acid assay (BCA) was used to quantify protein concentration. Between 5 and 15 μg protein input was used for WB experiments. Between 500 μg and 1000 μg protein input was used for pulldown for MS and the amount was adjusted to roughly normalize biotinylation levels across replicates. 10 μl of streptavidin magnetic beads (Pierce) per 100 μg of protein input was used for every condition. Beads were washed 3x in 0.5% SDS RIPA buffer using a magnetic rack and incubated overnight at 4 °C with rotation with protein input in 1 ml of 0.5% SDS RIPA buffer final volume. The following day, beads were washed 3x with 0.5% SDS RIPA buffer, 1x with 1 M KCl, 1x with 0.1 M Na2CO3, 3x with 2 M Urea in 50 mM Tris-HCl pH 8. At this point, proteins are ready for elution and MS sample preparation (described below).

### Western blot

After boiling 10 min at 95 °C, samples were loaded on 4–20% or 7.5% polyacrylamide gels and run at 180 V. Proteins were transferred to 0.2 um nitrocellulose membrane using semi-dry transfer (Biorad) using the mixed-molecular weight program. Immediately after transfer, a total protein stain (LI-COR Revert Total Protein Stain) was used to normalize loading. Then, membranes were blocked in 10% milk in TBS-T buffer (25 mM Tris-base pH 7.5, 300 mM NaCl, 0.05% Tween-20) for 30 mins at RT. Primary and secondary antibodies were diluted in 5% milk in TBS-T. Primary antibodies were incubated O/N at 4 °C, secondary antibodies were incubated 2 to 3 h at RT. When blotting for biotinylated proteins, membranes were thoroughly washed in TBS-T and HRP-conjugated streptavidin was diluted 1:500 in 5% BSA in TBS-T to avoid binding to the biotin present in the milk. After primary and secondary antibody incubation, membranes were washed 5 times with TBS-T and once with 1X PBS before development at ImageQuant 800 (Cytiva) using SuperSignal West Pico PLUS and Femto Chemiluminescent Substrate (Thermo Fisher 34577 and 34094).

### MS sample preparation

Proteins were eluted from the beads and digested to peptides using the micro S-Trap kit (Protifi) following manufacturer instructions. Briefly, beads were resuspended in 92 μl of 5% SDS 50 mM TEAB pH 8, boiled 5 min at 95 °C and let to cool down. Eluates were reduced with 4 μl of a 120 mM TCEP buffer at 55 °C for 20 mins, then alkylated with 4 μl of 500 mM MMTS in isopropanol at RT for 20 mins in the dark. Eluates were acidified with 24 μl of 27.5% phosphoric acid in dH2O. Using a magnetic rack to separate the beads, supernatants were collected in a different tube and 660 μl of binding buffer was added (100 mM TEAB in 90% methanol). Proteins were then loaded onto an S-trap column and centrifuged at 4000 G 1 min. Columns were washed 6 times with binding buffer. Proteins were digested by applying 1 μg of trypsin diluted in 20 μl of digestion buffer (50 mM TEAB) directly to the column and incubating overnight at 37 °C in a humidified oven. Peptides were sequentially eluted from the column using 40 μl of elution buffer 1, 2 and 3 and dried using a SpeedVac before being shipped for LC-MS/MS.

### Mass spectrometry

Peptide samples were analysed using an Evosep One liquid chromatography system (Evosep Biosystems) coupled online to an Orbitrap Astral mass spectrometer (Thermo Fisher Scientific) equipped with an EASY-Spray™ ion source. Peptides were loaded onto Evotips according to the manufacturer’s instructions and separated on an IonOpticks Aurora Elite TS column (15 cmx75 μm maintained at 50 °C) using the Whisper Zoom 40 samples per day (40SPD) method. For proteomics analyses, samples were resuspended in 20 µL of Buffer A (H_2_O supplemented with 0.1%FA). 1 µL of each sample was loaded.

The spray voltage was set to 1600 V, the funnel RF level to 40%, and the heated ion transfer tube at 300°C. The Orbitrap Astral mass spectrometer was operated in data-independent acquisition (DIA) mode at a full resolution of 240,000 over an m/z range of 380–980, with a normalized AGC target of 500% and a maximum injection time of 10 ms. DIA MS2 spectra were acquired in the Astral analyzer at a resolution of 80,000, using 2 Th isolation windows, a maximum injection time of 3 ms, and higher-energy collisional dissociation (HCD) at 25% normalized collision energy (NCE).

For the experiment described in Figure 5, 1 μl of the tryptic peptides were loaded onto a Vanquish Neo UHPLC system (Thermo Fisher Scientific) coupled to an Orbitrap Astral mass spectrometer (Thermo Fisher Scientific) operating in DIA mode. Solvent A consisted of 0.1% formic acid (FA) in water, and solvent B consisted of 80% acetonitrile (ACN) with 0.1% FA. Peptides were separated on an Aurora series column (TS version) 25 cm x 75 μm ID, 1.6 μm C18 (IonOpticks) maintained at 50°C. For this pilot study, a direct injection method was used with a total run time of 41.5 min. The gradient of solvent B (80% ACN, 0.1% FA) was as follows: 1–4% from 0–1.4 min at 550 nL/min; the flow rate was then reduced to 300 nL/min from 1.4–1.5 min while increasing solvent B from 4– 8%; subsequently, solvent B was increased to 31% from 1.5–26.5 min, to 45% from 26.5–31.5 min, and finally to 99%, which was held constant from 32.5–41.5 min. For Astral settings, MS1 spectra were acquired in the Orbitrap at a resolution of 240,000 over an m/z range of 380–980 with a normalized automatic gain control (AGC) target of 500%. DIA MS2 spectra were acquired in the Astral analyzer using a 0.6 s cycle time, 2 Th isolation windows, an AGC target of 500%, a maximum injection time of 3 ms, and a normalized collision energy of 25%.

### MS data analysis

For all experiments, RAW MS data were analysed with DIA-NN version 2.2.0, except for the experiment described in Figure 5 which used version 1.9.2. Uniprot *Mus musculus* UP000000589 reference proteome was used for spectral library preparation. Enzyme specificity was set to trypsin with up to one missed cleavage. Methionine oxidation and protein N-terminal acetylation were defined as variable modifications, while carbamidomethylation of cysteine was set as a fixed modification with a maximum of one occurrence. For DIA-NN 2.2.0, mass accuracy was set to 10 ppm and MS1 accuracy to 4.0 with the precursor FDR controlled at 1%. Matching between runs (MBR) was enabled, and scoring was set to peptidoforms. All other parameters were kept at default settings. For DIA-NN 1.9.2, the default settings were kept, using the first run to set mass and MS1 accuracies, and MBR was enabled

The unique.gene result file was loaded onto Perseus version 1.6 for analysis (Tyanova *et al*., 2016). Samples were log_2_ transformed. Only proteins present in all replicates in at least one condition were kept and missing values imputed with standard parameters. To enrich for synaptic proteins, we applied an ANOVA with Benjamini-Hochberg FDR followed by Tukey’s post-hoc HSD test (FDR=0.05) between the synaptic intrabody-TurboID and the Ctrl-turboID conditions, or a permutation-based t-Test with an FDR of 0.1 between Gphn-TurboID and Ctrl-TurboID in Fig. 5. We then merged significant proteins from all synaptic conditions when applicable (i.e. two synaptic probes in Fig. 2, three synaptic laminae in Fig. 3, or two timepoints and two mice strains in Fig. 6) to create the ‘synaptic protein list’ used for further comparison. For the synaptic laminae and AD datasets, this list was further filtered by cross-referencing with the Allen Brain 10X Dataset (parameters: CA1 Glut1-5 clusters, mean_exp>0.1, percent >0.2). To identify differentially expressed proteins between PSD95-TurboID and Homer1-TurboID (Fig. 2H), a permutation-based t-Test was performed on the synaptic proteins using FDR = 0.05 and s0 = 0.1 to calculate significance. To identify differentially expressed proteins in the synaptic laminae and AD datasets, we performed an ANOVA with Benjamini-Hochberg FDR followed by Tukey’s post-hoc HSD (FDR=0.05) to obtain a list of DEPs between laminae, or between AD and WT mice at each timepoint. From Perseus, results tables were exported as “.csv” for further analysis or data visualization using R.

### Protein ID numbers, Intensity plots and PCA analysis

Graphs showing protein ID numbers and intensity plots were obtained using DIA-analyst (https://analyst-suites.org/apps/dia-analyst/). PCA was performed after missing data imputation with standard parameters in Perseus.

### Cross-referencing with the Allen Brain Cell Atlas

Single-cell RNA-sequencing (scRNA-seq) data for the hippocampal formation (HPF) region were obtained from the Allen Brain Cell (ABC) Atlas (Yao *et al*., 2023). We downloaded the HPF datasets generated with 10x Genomics chemistries WMB-10xv2 and WMB-10xv3, retrieving the raw count matrices and the corresponding cell-level metadata. All quality control was performed in Scanpy (Wolf, Angerer and Theis, 2018). For each dataset (10Xv2 and 10Xv3) separately, we removed cells with less than 200 detected genes and genes detected in less than 3 cells. Percentage of mitochondrial genes was quantified per cell, and quality control metrics were computed with calculate_qc_metrics function; cells with more than 5% mitochondrial reads were excluded. Cells lacking a class-level annotation were also removed. In addition, we excluded mitochondrial and sex-linked genes including: *Ehd2, Xist, Tsix, Eif2s3y, Ddx3y, Uty, Kdm5d, Gstp1, Rpl35a, Erh, Slc25a5, Pgk1, Eno1, Tubb2a, Emc4, Scg5,* and *Rn45s* (as referenced in (Rasetto *et al*., 2024)). The WMB-10Xv2 and WMB-10Xv3 datasets were merged. Glutamatergic neurons were then isolated, and genes detected in fewer than three cells were removed. CA1 cells were then extracted by selecting cells belonging to CA1 supertypes Glut1–Glut5 (excluding Glut6). Counts were normalized to 10,000 UMIs per cell with normalize_total function and subsequently log transformed. For our predefined list of genes of interest, expression metrics were computed from the log-normalized counts by computing (i) mean expression (average across cells) and (ii) percent expressing cells (fraction of cells with nonzero expression). To guide cutoff selection, histograms of mean expression and detection rate were generated with matplotlib (Hunter, 2007). Genes with mean expression higher than 0.1 and percent expression higher than 0.2 were retained for downstream analyses.

### Gene ontology analysis and protein annotation

GO term enrichment was done in Perseus (annotated volcano plots or imported to R and analyzed with the packages clusterProfiler and mSigdb (Subramanian *et al*., 2005; Yu *et al*., 2012; Liberzon *et al*., 2015). SynGO enrichment was performed using SynGO’s webtool (Koopmans *et al*., 2019)(syngoportal.org). The lists of E3 ligases and RNA binding proteins were obtained from (Medvar *et al*., 2016) and (Gerstberger, Hafner and Tuschl, 2014), respectively.

## Data availability

RAW MS files will be deposited on public servers such as Proteomexchange or PRIDE upon submission. Any additional information required to reanalyze the data reported in this paper is available from the lead contact upon request.

## Acknowledgements

We thank Keimpe Wierda, Patrik Verstreken, Esther Klingler and Bart De Strooper for critical reading of the manuscript, and De Wit lab members for helpful discussion and comments. We thank ThermoFisher Scientific for demonstrating the Astral Mass Spectrometer and providing the data in Figure 5E. We are grateful to Joris Vandenbempt for mouse colony maintenance and genotyping. We thank Marinka Brouwer, Stephanie Chanda and Beatriz Marques for help with primary neuronal cultures. D.D. is supported by Fonds Wetenschappelijk Onderzoek (FWO) Postdoctoral fellowship 12W5218N, 12W5221N and FWO Research Grant 1513320N; G.M. is supported by FWO PhD fellowship 11F1219N and 11F1221N, and the Fund Janine et Jacques Delruelle; B.L-E. is supported by FWO PhD fellowship 1120821 N and 1120823 N; A.K. is supported by a Marie Skłodowska-Curie Actions Post-doctoral fellowship HORIZON-MSCA-2024-PF-01 Project 101201966 and a Fund Doctor Gustave Delport (managed by King Baudouin Foundation) Project 2024-J1811190-0023070; J.d.W. is supported by FWO Project Grants G0A8720N, G0A8320N, G0ABV25N and G0AJM25N; FWO-SBO S005024N Enhancer-AI; the Queen Elisabeth Medical Foundation for Neurosciences; Stichting Alzheimer Onderzoek-Fondation Recherche Alzheimer (SAO-FRA) Standard Grant 20230028, and a Methusalem Grant of KU Leuven/Flemish Government. This investigation was also supported by the FWO Large-scale Research Infrastructure Grant I015524N.

## Notes

### Competing Interest Statement

J.d.W. is scientific co-founder and served as scientific advisory board member of Augustine Tx. The remaining authors declare no competing interests.

