## Supplementary Figures for "Intrabody-guided synapse proteomics defines pyramidal neuron input architecture and uncovers early remodeling in a mouse model of Alzheimer’s disease"

### Figure S1

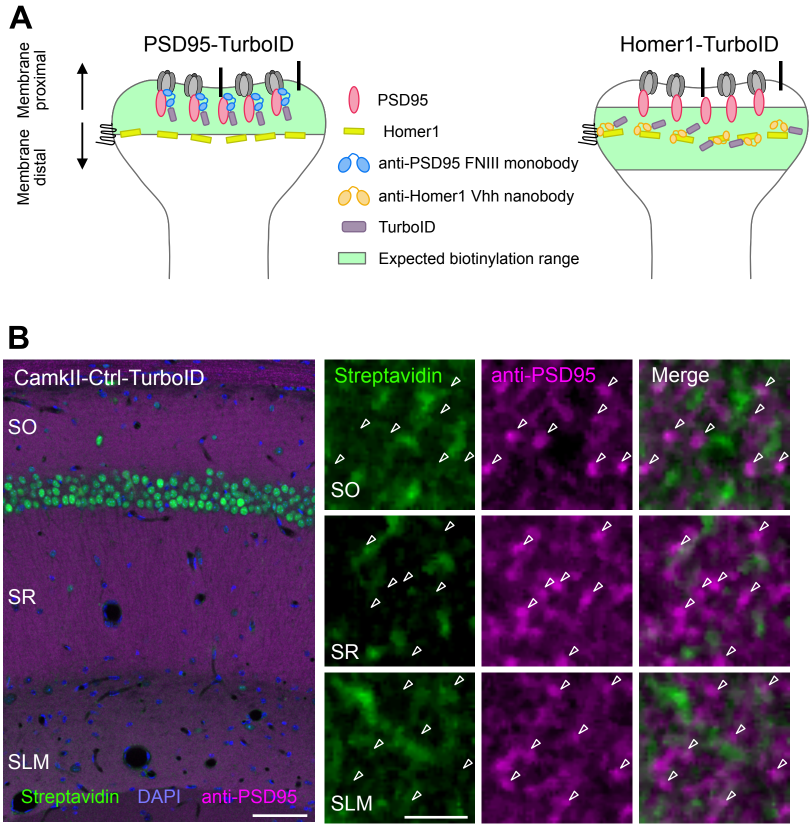
A
PSD95-TurboID
Homer1-TurboID
Membrane proximal
Membrane distal
PSD95
Homer1
anti-PSD95 FNIII monobody
anti-Homer1 Vhh nanobody
TurboID
Expected biotinylation range
B
CamkII-Ctrl-TurboID
SO
SR
SLM
Streptavidin
DAPI
anti-PSD95
Merge

#### Figure S2

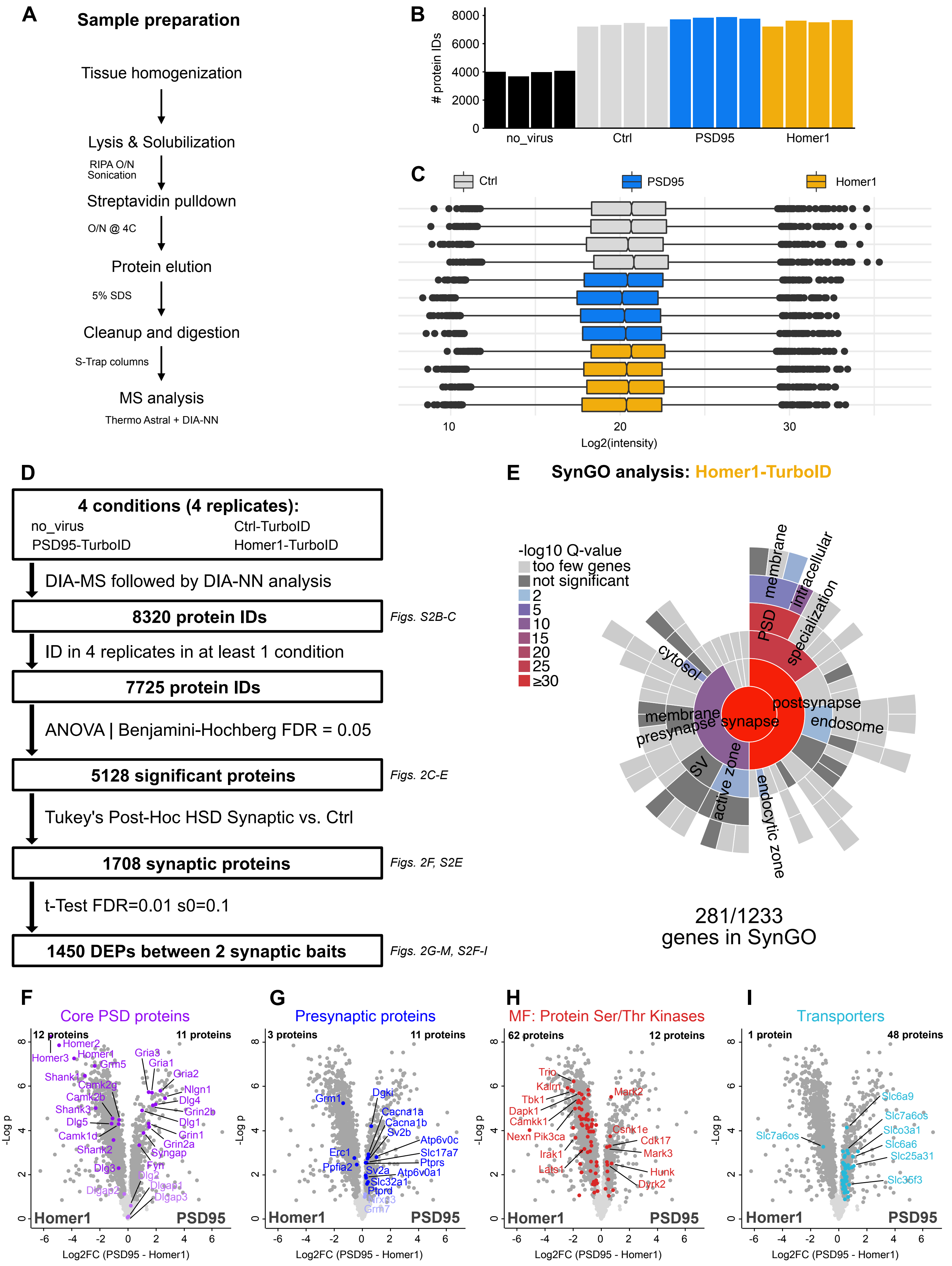
A
Sample preparation
Tissue homogenization
Lysis & Solubilization
RIPA O/N
Sonication
Streptavidin pulldown
O/N @ 4C
Protein elution
5% SDS
Cleanup and digestion
S-Trap columns
MS analysis
Thermo Astral + DIA-NN
B
### protein IDs
no_virus
Ctrl
PSD95
Homer1
C
Ctrl
PSD95
Homer1
Log2(intensity)
D
4 conditions (4 replicates):
no_virus
Ctrl-TurboID
PSD95-TurboID
Homer1-TurboID
DIA-MS followed by DIA-NN analysis
8320 protein IDs
Figs. S2B-C
ID in 4 replicates in at least 1 condition
7725 protein IDs
ANOVA | Benjamini-Hochberg FDR = 0.05
5128 significant proteins
Figs. 2C-E
Tukey's Post-Hoc HSD Synaptic vs. Ctrl
1708 synaptic proteins
Figs. 2F, S2E
t-Test FDR=0.01 s0=0.1
1450 DEPs between 2 synaptic baits
Figs. 2G-M, S2F-I
E
SynGO analysis: Homer1-TurboID
-log10 Q-value
too few genes
not significant
2
5
10
15
20
25
≥30
membrane
intracellular
PSD
specialization
cytosol
postsynapse
endosome
membrane
presynapse
synapse
SV
active zone
endocytic zone
281/1233
genes in SynGO
F
Core PSD proteins
12 proteins
11 proteins
Homer1
PSD95
-Log p
Log2FC (PSD95 - Homer1)
G
Presynaptic proteins
3 proteins
11 proteins
H
MF: Protein Ser/Thr Kinases
62 proteins
12 proteins
I
Transporters
1 protein
48 proteins

Figure S3

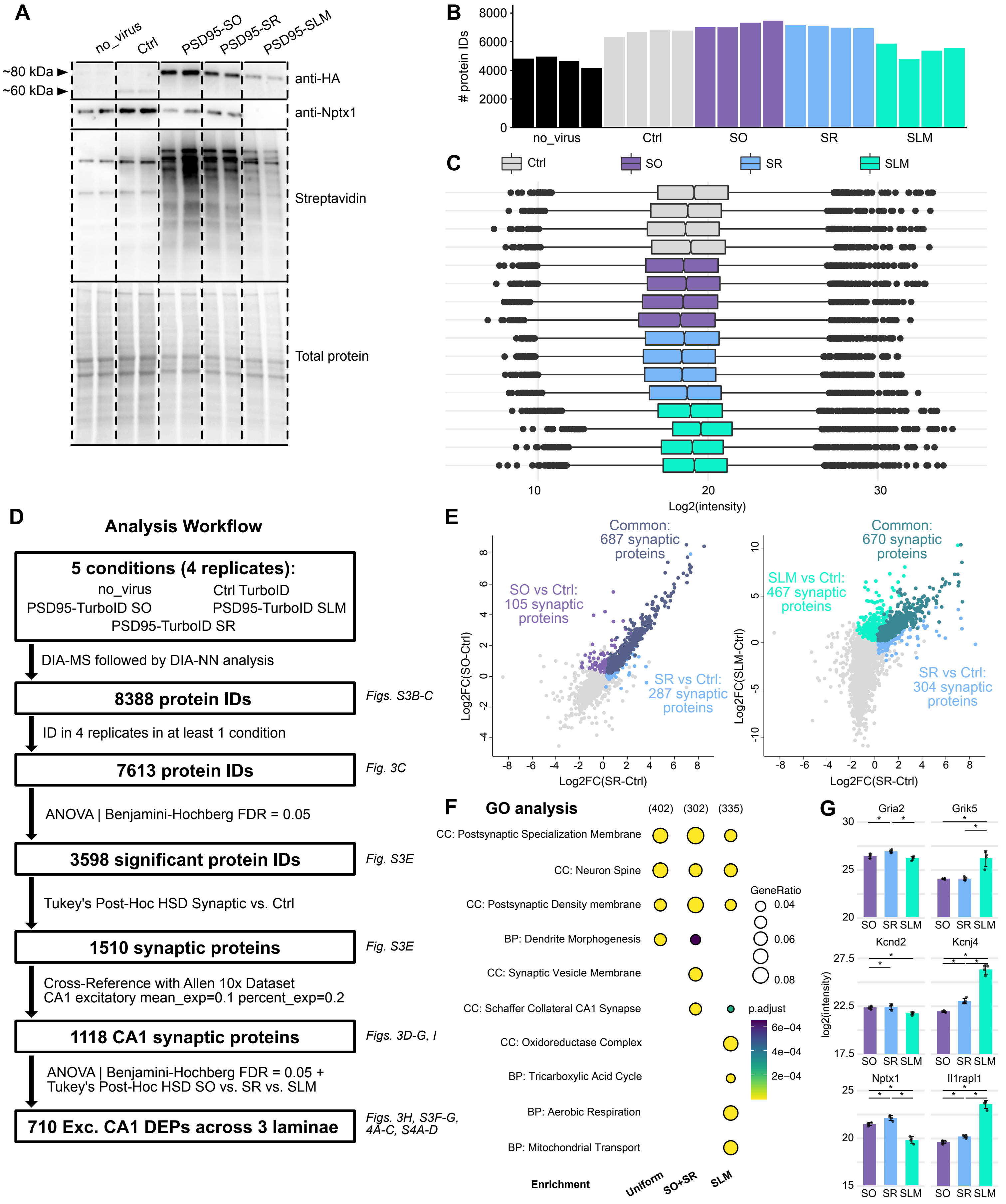
A
no_virus
Ctrl
PSD95-SO
PSD95-SR
PSD95-SLM
~80 kDa ►
~60 kDa ►
anti-HA
anti-Nptx1
Streptavidin
Total protein
B
### protein IDs
8000
6000
4000
2000
0
no_virus
Ctrl
SO
SR
SLM
C
Ctrl
SO
SR
SLM
10
20
30
Log2(intensity)
D
Analysis Workflow
5 conditions (4 replicates):
no_virus
Ctrl TurboID
PSD95-TurboID SO
PSD95-TurboID SLM
PSD95-TurboID SR
DIA-MS followed by DIA-NN analysis
8388 protein IDs
Figs. S3B-C
ID in 4 replicates in at least 1 condition
7613 protein IDs
Fig. 3C
ANOVA | Benjamini-Hochberg FDR = 0.05
3598 significant protein IDs
Fig. S3E
Tukey's Post-Hoc HSD Synaptic vs. Ctrl
1510 synaptic proteins
Fig. S3E
Cross-Reference with Allen 10x Dataset
CA1 excitatory mean_exp=0.1 percent_exp=0.2
1118 CA1 synaptic proteins
Figs. 3D-G, I
ANOVA | Benjamini-Hochberg FDR = 0.05 +
Tukey's Post-Hoc HSD SO vs. SR vs. SLM
710 Exc. CA1 DEPs across 3 laminae
Figs. 3H, S3F-G,
4A-C, S4A-D
E
Common:
687 synaptic
proteins
SO vs Ctrl:
105 synaptic
proteins
SR vs Ctrl:
287 synaptic
proteins
Log2FC(SO-Ctrl)
Log2FC(SR-Ctrl)
Common:
670 synaptic
proteins
SLM vs Ctrl:
467 synaptic
proteins
SR vs Ctrl:
304 synaptic
proteins
Log2FC(SLM-Ctrl)
Log2FC(SR-Ctrl)
F
GO analysis
(402)
(302)
(335)
CC: Postsynaptic Specialization Membrane
CC: Neuron Spine
CC: Postsynaptic Density membrane
BP: Dendrite Morphogenesis
CC: Synaptic Vesicle Membrane
CC: Schaffer Collateral CA1 Synapse
CC: Oxidoreductase Complex
BP: Tricarboxylic Acid Cycle
BP: Aerobic Respiration
BP: Mitochondrial Transport
GeneRatio
0.04
0.06
0.08
p.adjust
6e−04
4e−04
2e−04
Enrichment
Uniform
SO+SR
SLM
G
Gria2
Grik5
Kcnd2
Kcnj4
Nptx1
Il1rapl1
log2(intensity)
SO
SR
SLM

#### Figure S4

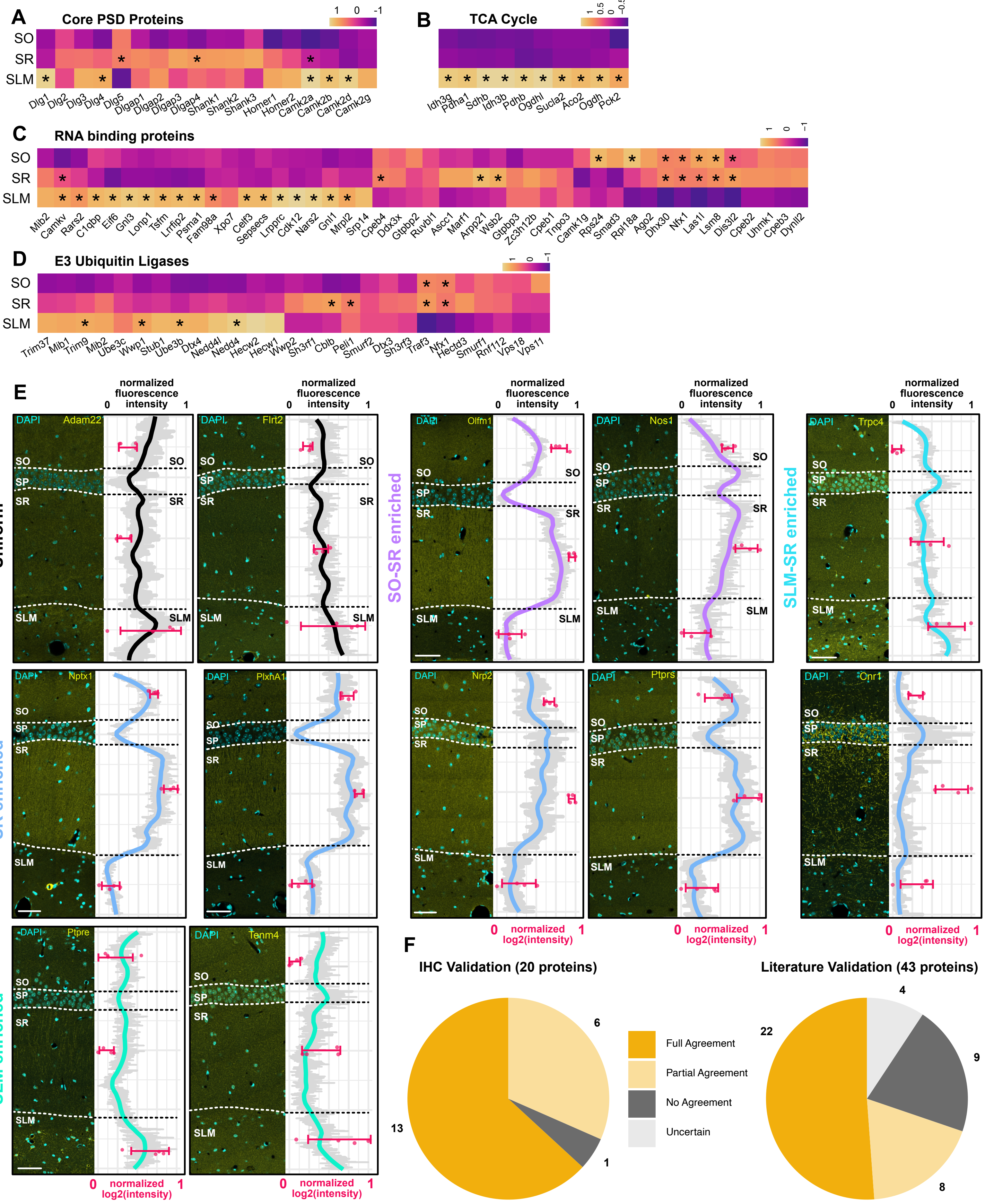

A
Core PSD Proteins
SO
SR
SLM
Dlg1 Dlg2 Dlg3 Dlg4 Dlg5 Dlgap1 Dlgap2 Dlgap3 Dlgap4 Shank1 Shank2 Shank3 Homer1 Homer2 Camk2a Camk2b Camk2d Camk2g
B
TCA Cycle
Idh3g Pdha1 Sdhb Idh3b Pdhb Ogdhl Sucla2 Aco2 Ogdh Pck2
C
RNA binding proteins
Mib2 Camkv Rars2 C1qbp Eif6 Gnl3 Lonp1 Tsfm Lrrfip2 Psma1 Fam98a Xpo7 Celf3 Sepsecs Lrpprc Cdk12 Nars2 Gnl1 Mrpl2 Srp14 Cpeb4 Ddx3x Gtpbp2 Ruvbl1 Ascc1 Marf1 Arpp21 Wsb2 Gtpbp3 Zc3h12b Cpeb1 Tnpo3 Camk1g Rps24 Smad3 Rpl18a Ago2 Dhx30 Nfx1 Las1l Lsm8 Dis3l2 Cpeb2 Unmk1 Cpeb3 Dynll2
D
E3 Ubiquitin Ligases
Trim37 Mib1 Trim9 Mib2 Ube3c Wwp1 Stub1 Ube3b Dtx4 Nedd4l Nedd4 Hecw2 Hecw1 Wwp2 Sh3rf1 Cblb Peli1 Smurf2 Dtx3 Sh3rf3 Traf3 Nfx1 Hectd3 Smurf1 Rnf112 Vps18 Vps11
E
normalized fluorescence intensity
Uniform
SO-SR enriched
SLM-SR enriched
SR enriched
SLM enriched
DAPI Adam22
DAPI Flrt2
DAPI Olfm1
DAPI Nos1
DAPI Trpc4
DAPI Nptx1
DAPI PlxnA1
DAPI Nrp2
DAPI Ptprs
DAPI Cnr1
DAPI Ptpre
DAPI Tenm4
SO SP SR SLM
normalized log2(intensity)
F
IHC Validation (20 proteins)
Literature Validation (43 proteins)
Full Agreement
Partial Agreement
No Agreement
Uncertain
13 6 1
22 4 9 8

**Figure S5**

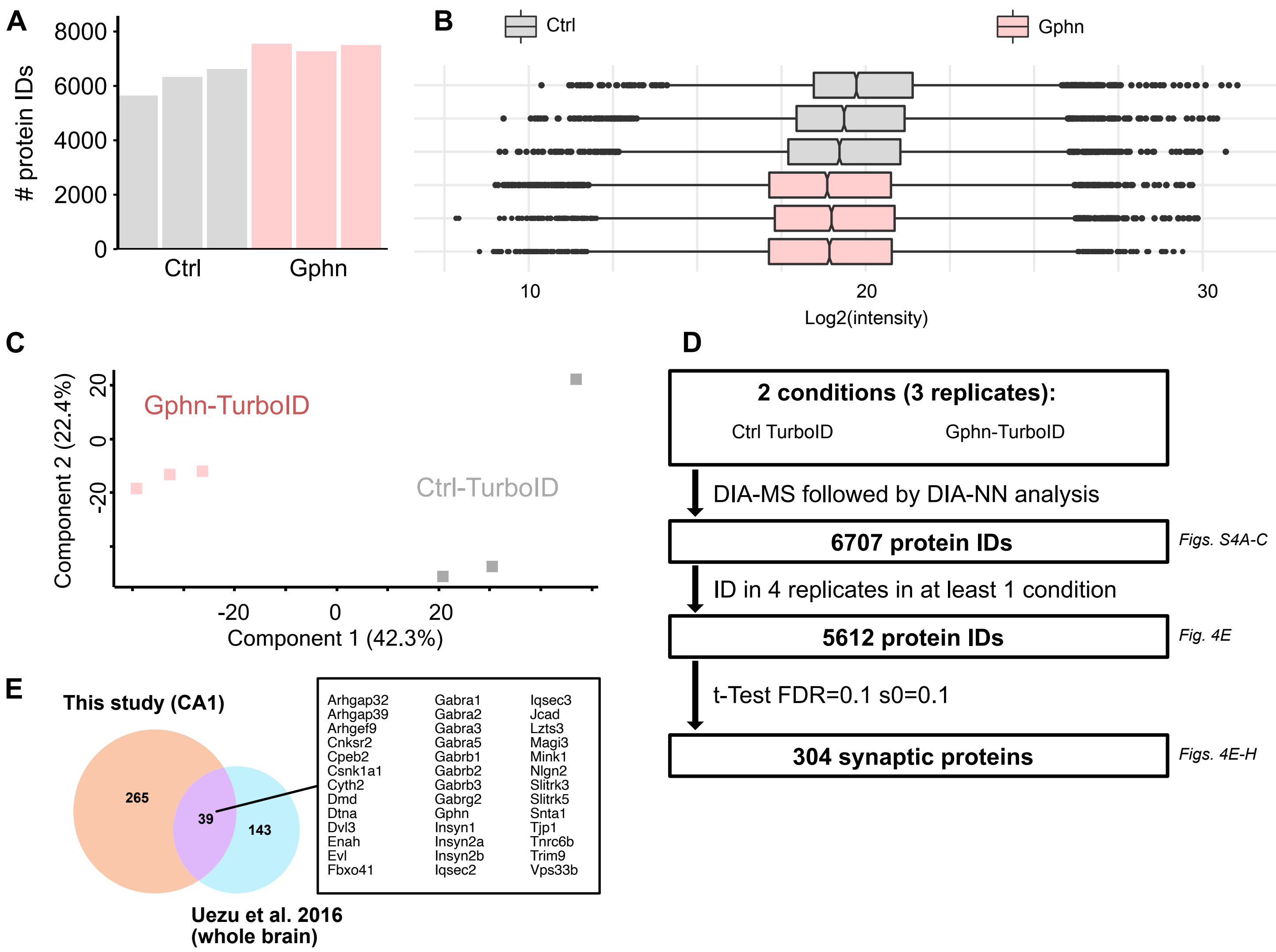
A
8000
6000
4000
2000
0
### protein IDs
Ctrl
Gphn
B
Ctrl
Gphn
10
20
30
Log2(intensity)
C
Gphn-TurboID
Ctrl-TurboID
Component 2 (22.4%)
20
0
-20
-20
0
20
Component 1 (42.3%)
D
2 conditions (3 replicates):
Ctrl TurboID
Gphn-TurboID
DIA-MS followed by DIA-NN analysis
6707 protein IDs
Figs. S4A-C
ID in 4 replicates in at least 1 condition
5612 protein IDs
Fig. 4E
t-Test FDR=0.1 s0=0.1
304 synaptic proteins
Figs. 4E-H
E
This study (CA1)
265
39
143
Uezu et al. 2016
(whole brain)
Arhgap32
Arhgap39
Arhgef9
Cnksr2
Cpeb2
Csnk1a1
Cyth2
Dmd
Dtna
Dvl3
Enah
Evl
Fbxo41
Gabra1
Gabra2
Gabra3
Gabra5
Gabrb1
Gabrb2
Gabrb3
Gabrg2
Gphn
Insyn1
Insyn2a
Insyn2b
Iqsec2
Iqsec3
Jcad
Lzts3
Magi3
Mink1
Nlgn2
Slitrk3
Slitrk5
Snta1
Tjp1
Tnrc6b
Trim9
Vps33b

#### Figure S6

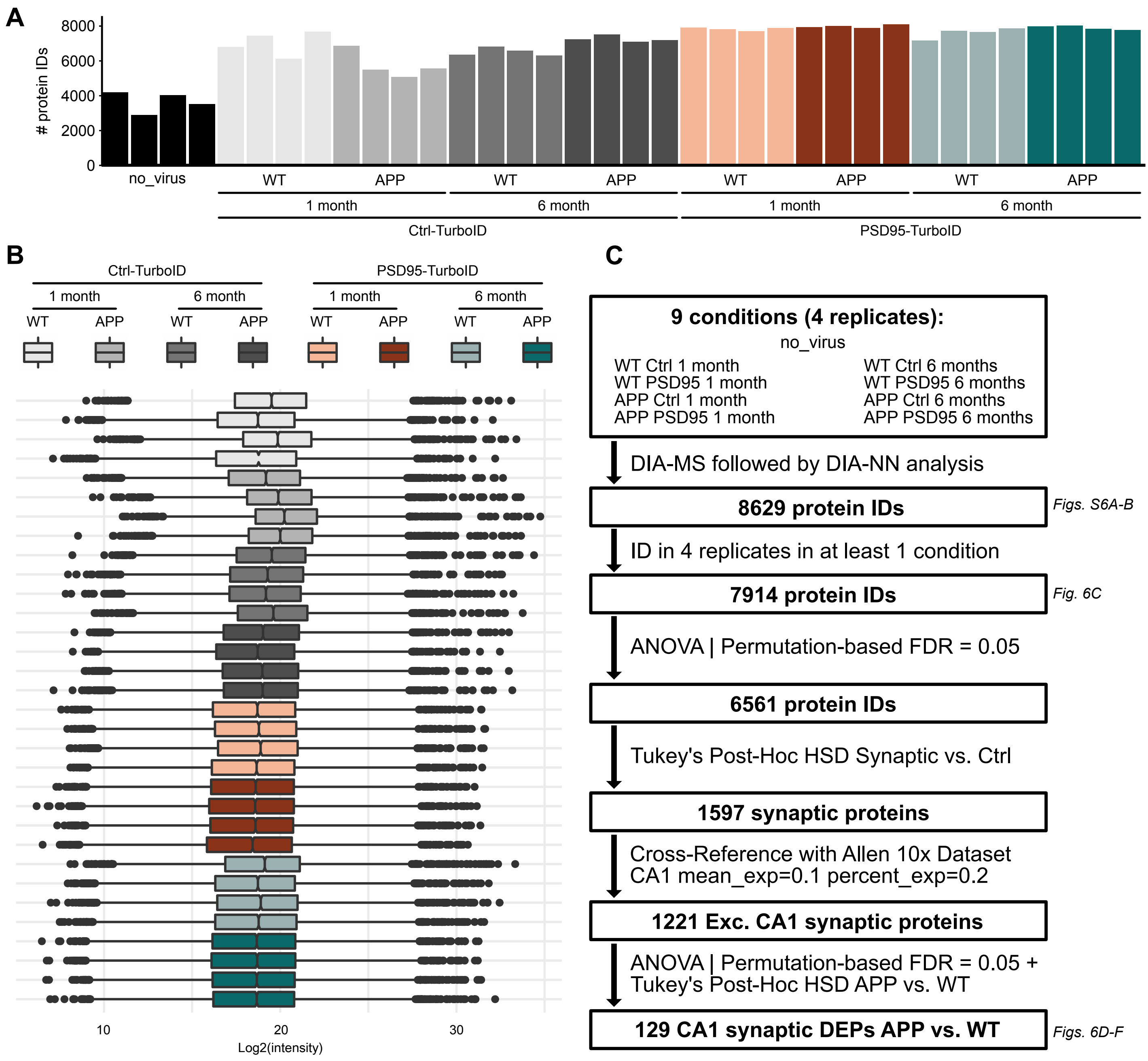
A
### protein IDs
8000
6000
4000
2000
0
no_virus
WT
APP
WT
APP
WT
APP
WT
APP
1 month
6 month
1 month
6 month
Ctrl-TurboID
PSD95-TurboID
B
Ctrl-TurboID
PSD95-TurboID
1 month
6 month
1 month
6 month
WT
APP
WT
APP
WT
APP
WT
APP
10
20
30
Log2(intensity)
C
9 conditions (4 replicates):
no_virus
WT Ctrl 1 month
WT PSD95 1 month
APP Ctrl 1 month
APP PSD95 1 month
WT Ctrl 6 months
WT PSD95 6 months
APP Ctrl 6 months
APP PSD95 6 months
DIA-MS followed by DIA-NN analysis
8629 protein IDs
Figs. S6A-B
ID in 4 replicates in at least 1 condition
7914 protein IDs
Fig. 6C
ANOVA | Permutation-based FDR = 0.05
6561 protein IDs
Tukey's Post-Hoc HSD Synaptic vs. Ctrl
1597 synaptic proteins
Cross-Reference with Allen 10x Dataset
CA1 mean_exp=0.1 percent_exp=0.2
1221 Exc. CA1 synaptic proteins
ANOVA | Permutation-based FDR = 0.05 +
Tukey's Post-Hoc HSD APP vs. WT
129 CA1 synaptic DEPs APP vs. WT
Figs. 6D-F

#### Figure S7

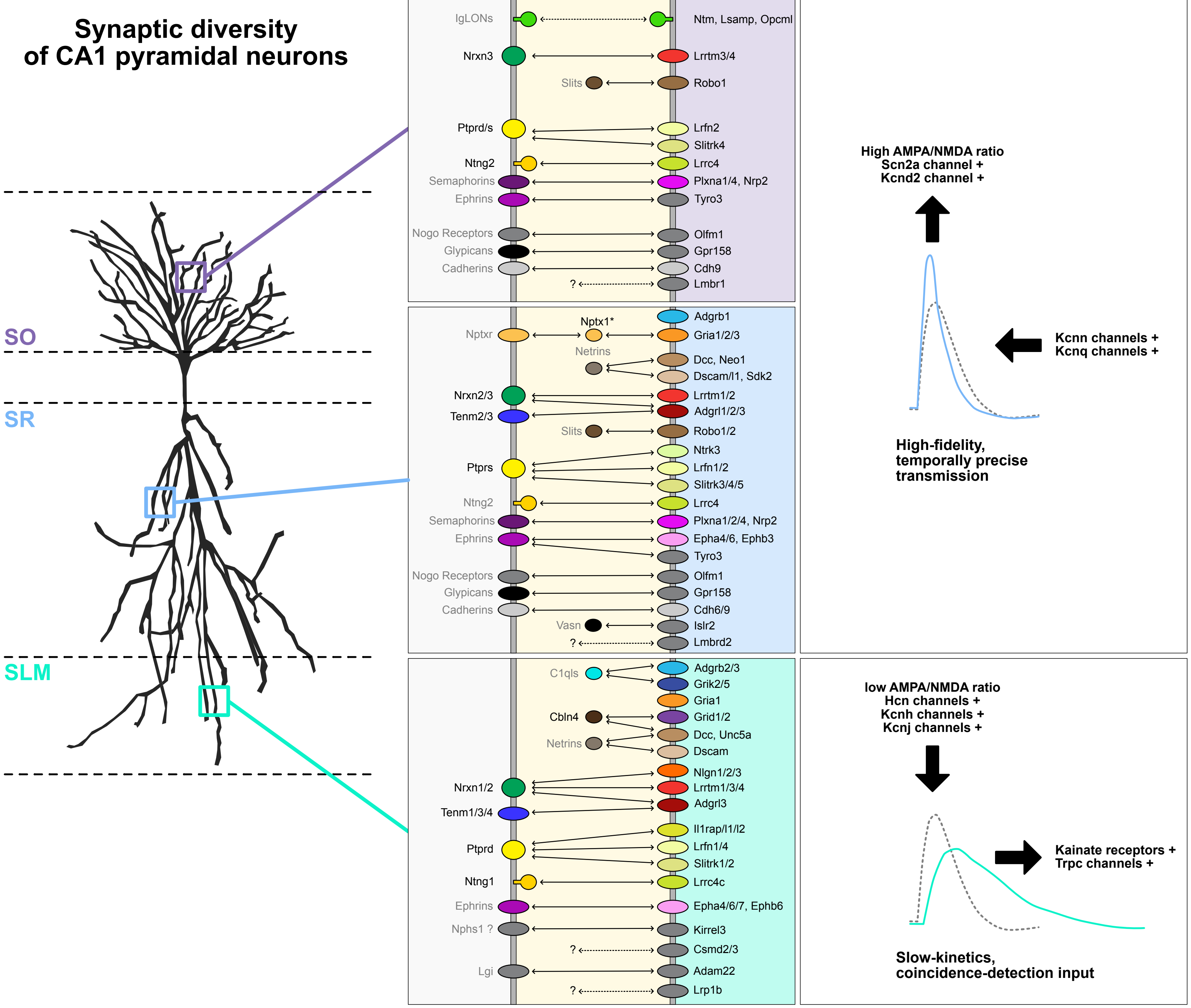
Synaptic diversity
of CA1 pyramidal neurons
SO
SR
SLM
Nptx1
Nptxr
Gria4
IgLONs
Ntm, Lsamp, Opcml
Nrxn3
Lrrtm3/4
Slits
Robo1
Ptprd/s
Lrfn2
Slitrk4
Ntng2
Lrrc4
Semaphorins
Plxna1/4, Nrp2
Ephrins
Tyro3
Nogo Receptors
Olfm1
Glypicans
Gpr158
Cadherins
Cdh9
?
Lmbr1
Adgrb1
Nptx1*
Nptxr
Gria1/2/3
Netrins
Dcc, Neo1
Dscam/l1, Sdk2
Nrxn2/3
Lrrtm1/2
Tenm2/3
Adgrl1/2/3
Slits
Robo1/2
Ntrk3
Ptprs
Lrfn1/2
Slitrk3/4/5
Ntng2
Lrrc4
Semaphorins
Plxna1/2/4, Nrp2
Ephrins
Epha4/6, Ephb3
Tyro3
Nogo Receptors
Olfm1
Glypicans
Gpr158
Cadherins
Cdh6/9
Vasn
Islr2
?
Lmbrd2
C1qls
Adgrb2/3
Grik2/5
Gria1
Cbln4
Grid1/2
Netrins
Dcc, Unc5a
Dscam
Nlgn1/2/3
Nrxn1/2
Lrrtm1/3/4
Adgrl3
Tenm1/3/4
Il1rap/l1/l2
Ptprd
Lrfn1/4
Slitrk1/2
Ntng1
Lrrc4c
Ephrins
Epha4/6/7, Ephb6
Nphs1 ?
Kirrel3
?
Csmd2/3
Lgi
Adam22
?
Lrp1b
High AMPA/NMDA ratio
Scn2a channel +
Kcnd2 channel +
Kcnn channels +
Kcnq channels +
High-fidelity,
temporally precise
transmission
low AMPA/NMDA ratio
Hcn channels +
Kcnh channels +
Kcnj channels +
Kainate receptors +
Trpc channels +
Slow-kinetics,
coincidence-detection input

**Supplementary figure legends**

**Figure S1.** A. Different sub-synaptic proteomes expected to be enriched in the PSD95-TurboID and Homer1-TurboID conditions. B. Left: Low-magnification z-stacks of dorsal CA1 in animals injected with Ctrl-TurboID. Right: High-magnification single planes showing no synaptic biotinylation *in vivo* in the three CA1 synaptic laminae. Scale bars are: 100 μm (B left), 2 μm (B right).

**Figure S2.** A. Simplified representation of the sample preparation and MS workflow. B. Number of protein IDs in each sample. C. Distribution of MS intensities in each sample. D. Data analysis workflow. E. SynGO enrichment diagram for the Homer1-TurboID condition. F to I. Differential enrichment of different protein groups in the Homer1-TurboID or PSD95-TurboID conditions.

**Figure S3**. A. Western Blot showing expression of PSD95-TurboID and Ctrl-TurboID probes, as well as biotinylation levels for two out of four replicates in the experiment. Additionally, anti-Nptx1 staining shows the accuracy of the microdissection. B. Number of protein IDs in each sample. C. Distribution of MS intensities in each sample. D. Data analysis workflow. E. Plots showing the filtering against Ctrl-TurboID for the SO and SR condition (left) and the SLM and SR condition (right) (ANOVA with Benjamini-Hochberg FDR=0.05 followed by Tukey's HSD). F. GO analysis of Uniform, SO+SR, and SLM enriched proteomes using GO:CC and GO:BP terms. Numbers in parenthesis indicate the number of annotated proteins in each group. G. Examples of raw MS intensity plots for a few proteins.

**Figure S4**. A to D. Heatmaps showing the relative distribution across CA1 synaptic laminae of PSD proteins (A), TCA cycle proteins (B), RNA binding proteins (C), and E3 Ubiquitin ligases (D). E. More examples of hippocampal CA1 stainings against different proteins in the laminar dataset. F. Pie charts illustrating the proportion of proteins with matching distribution using IHC or datamining. Abbreviations: SO: *stratum oriens*, SR: *stratum radiatum*, SLM: *stratum lacunosum-moleculare*.

**Figure S5**. A. Number of protein IDs in each sample. B. Distribution of MS intensities in each sample. C. PCA showing the separation of the two experimental conditions based on MS intensities. D. Data analysis workflow. E. Comparison with a previous study using proximity biotinylation in inhibitory synapses of the whole brain.

**Figure S6.** A. Number of protein IDs in each sample. B. Distribution of MS intensities in each sample. C. Data analysis workflow.

**Figure S7. Proposed model for the CA1 proximo-distal molecular logic.** Left: CSP distribution based on our results and published literature. In black are proteins present in our dataset (regardless of CA1 expression). In gray are proteins inferred from known ligand-receptor databases. Right: Model of the functional consequences of different molecular synaptic signatures. Figure design inspired by Südhof, 2018.
